# Thymidine kinase-independent click chemistry DNADetect™ probes as an EdU alternative for mammalian cell DNA labelling

**DOI:** 10.1101/2025.04.02.646904

**Authors:** Jacinta R. Macdonald, Gillian M. Fisher, Anne-Sophie C. Braun, Shilpa Thomas, Tina S. Skinner-Adams, David H. Hilko, Sally-Ann Poulsen, Katherine T. Andrews

## Abstract

Nucleoside analogues have been powerful tools to study DNA synthesis, cell cycle progression and cellular fate for decades. EdU (5-ethynyl-2′-deoxyuridine), which contains an alkyne handle that allows copper-catalyzed azide-alkyne cycloaddition (CuAAC) to a fluorescent azide in a “click” chemistry reaction, is one of the most frequently utilised nucleoside analogue probes. EdU is transported into cells by nucleoside transporters followed by phosphorylation by thymidine kinase, the first step in thymidine metabolism leading to DNA synthesis. As some organisms like malaria parasites lack the thymidine kinase enzyme and cannot be labelled by EdU or related thymidine analogue probes, we previously designed and validated DNADetect™ EdU analogues as chemical probes. These probes comprise a masked monophosphate on the 5′-hydroxyl group of the nucleoside sugar moiety that is metabolised directly to EdU monophosphate, bypassing the need for thymidine kinase. Here, we demonstrate that DNADetect™ probes can be used to label DNA in mammalian cells. DNADetect™ probes are incorporated into proliferating HeLa cells as efficiently as EdU and outcompeted by thymidine. Additionally, we implement a protocol for best practice use of metabolic chemical probes by using a specifically designed inactive control probe for each active probe. While this approach is commonly applied with chemical probes that modulate protein function, it is yet to be commonly applied with metabolic chemical probes.

## Introduction

Nucleoside analogues are used for a variety of applications including as chemical probes to investigate DNA synthesis, cell cycle progression and other cellular processes (reviewed in (Solius et al., 2021)). Methods of labelling have progressed from radiolabel-detection using ^3^H-thymidine to other imaging-based detection methods which utilise antibodies and/or chemical-based approaches to detect incorporated nucleosides (Solius et al., 2021). The 5-halo-pyrimidine nucleoside 5-bromo-2’-deoxyuridine, BrdU, was developed to detect DNA replication, with incorporation determined by autoradiography (Djordjevic & Szybalski, 1960; Eidinoff et al., 1959), and later adapted to detection with monoclonal antibodies (Gratzner, 1982; Sawicki et al., 1971). However, a limitation of BrdU is that the antibody epitope is sterically hindered by complimentary base pairing in double-stranded DNA and requires harsh denaturation conditions for antibody access (Marti-Clua, 2024; Solius et al., 2021). 5-Ethynyl-2′-deoxyuridine (EdU), an analogue of thymidine in which a 5-ethynyl group replaces the 5-methyl group, overcomes the need for harsh denaturing conditions by enabling detection via a covalent reaction with a fluorescent azide to the 5-ethynyl group with a Cu(I)-catalyzed [3 + 2] cycloaddition reaction, commonly described as a “click” chemistry reaction (Buck et al., 2008; Ligasova & Koberna, 2018; Solius et al., 2021). The fluorescent azide (MW <1 kDa) is substantially smaller than detection antibodies (MW ∼150 kDa), removing the need for DNA denaturation to access the modified thymidine. Depending on the probe and biological application, other considerations for the use of BrdU, EdU, and related nucleoside analogues include bioavailability, toxicity (e.g., for extended incubation times) and the requirement for active nucleoside transport into the cell and recognition and processing by a functional thymidine kinase, the first step in nucleoside metabolism leading to eventual DNA synthesis and probe incorporation (Bitter et al., 2020; Solius et al., 2021; Wright & Lee, 2021). Additionally, corresponding control probes that address the impact of all the chemical probe modifications as compared to the canonical endogenous ligand thymidine should be considered for validation of chemical probes in new biological systems.

Nucleosides and nucleoside analogues like EdU have minimal membrane permeability due to their hydrophilicity and instead are actively transported into the cells by two families of nucleoside transporters (NTs) (Wright & Lee, 2021). In human cells these are sodium-dependant concentrative nucleoside transporters (hCNTs) and sodium-independent equilibrative nucleoside transporters (hENTs) (Wright & Lee, 2021; Young et al., 2013). The functional mechanisms of these two transporter families are distinct with individual transporters demonstrating solute specific activity (Pastor-Anglada & Perez-Torras, 2018; Wright & Lee, 2021). In addition, the tissue and subcellular location of these transporters can vary, with each being regulated by multiple mechanisms (Pastor-Anglada & Perez-Torras, 2018; Wright & Lee, 2021; Young et al., 2013). Thus, variations in transporter location, expression, and function due to a disease state (e.g., tumour cell lines (Farre et al., 2004; Perez-Torras et al., 2013) or Crohn’s disease (Perez-Torras et al., 2016)), may impact DNA labelling studies with probes like EdU that are actively transported into cells, impacting both *in vitro* and *in vivo* use. Like nucleoside prodrugs that overcome the polarity of nucleosides and hence their dependence on transporters (e.g., NUC-1031 and NUC-3373, reviewed in (Xu et al., 2023); and tenofovir disoproxil fumarate (TDF), reviewed in (Pradere et al., 2014)), transporter-independent lipophilic EdU analogues are of interest as chemical probes for biolabeling applications, for example as we previously showed for phosphotriester pronucleotide EdU analogues (Huynh et al., 2015).

In recent work (Hilko et al., 2023), we designed and synthesized EdU analogues with an acyloxybenzyl protected phosphate group on the 5′-hydroxyl group of the nucleoside sugar moiety (DNADetect™ probes **1a-1d**; **Figure 1**), three of which have lipophilicity that is in the range consistent with good cellular uptake by passive membrane permeability (**1b-1d**; cLog P, 4.24, 3.94, and 5.40, respectively) and *in vitro* stability suitable for labelling assays in *Plasmodium falciparum* malaria parasites (half-life in media 1.60 h, 1.66 h and 0.53 h, respectively) (Hilko et al., 2023). These probes were designed to address the limitation that EdU cannot be utilised in *Plasmodium* parasites, which lack thymidine kinase, the first and rate-limiting enzyme in the successive phosphorylation steps needed to convert nucleosides to nucleotides for DNA synthesis (Bitter et al., 2020), and furthermore have multiple membrane barriers to reach the parasites nucleus. We demonstrated that DNADetect™ probes **1b-1d** label *P. falciparum* DNA (Hilko et al., 2023), providing an EdU alternative for this parasite and other organisms lacking thymidine kinase. In contrast, **1a**, which is predicted to be predominantly hydrolysed outside the cell (half-life in media 0.085 h) had poor labelling, consistent with *Plasmodium* lacking thymidine kinase (Hilko et al., 2023). In this study, we aimed to expand the scope of applications of DNADetect™ probes **1a-1d** as transporter-independent and thymidine kinase-independent alternatives to EdU for use in mammalian cells, investigating their cytotoxicity and ability to label proliferating cells using flow cytometry and fluorescence microscopy. Furthermore, we demonstrated that **1d** retains the ability to be outcompeted by thymidine, a trait shared by EdU (Ligasová et al., 2015).

**Figure 1.**
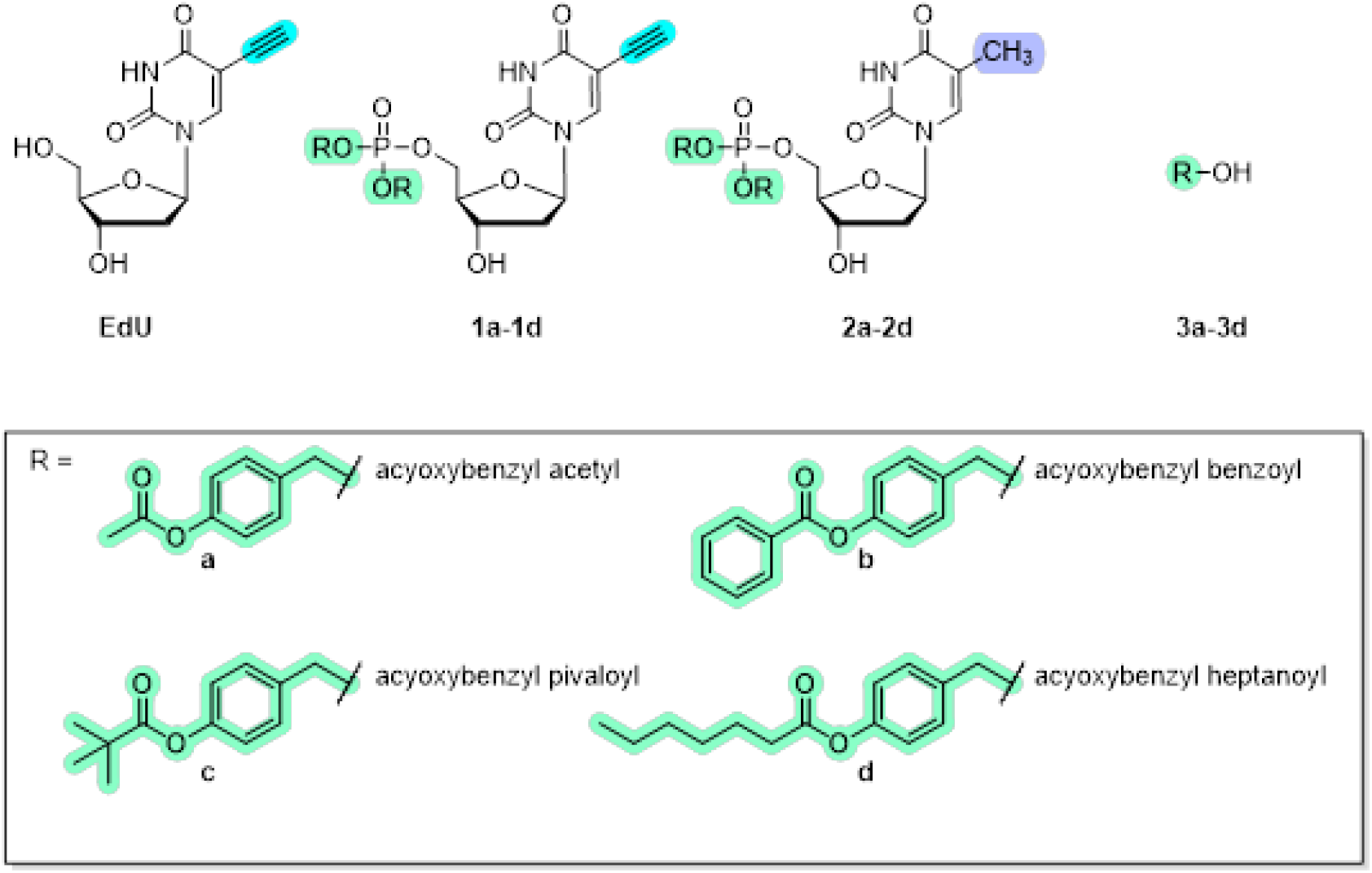
EdU, DNADetect™ probes **1a-1d**, controls **2a-2d**, and acyloxybenzyl alcohols **3a-3d**.

## Results and Discussion

### Cytotoxicity of DNADetect™ probes

To ensure that labelling concentration ranges were not toxic to replicating human cells, probes **1a-1d** were first assessed for *in vitro* growth inhibition of HeLa cells after 4 h and 24 h exposure using sulforhodamine B (SRB) assays (Skehan et al., 1990). Controls included the thymidine probes, **2a-2d** (identical to **1a-1d**, respectively, but retain the methyl group of thymidine instead of the alkyne group of EdU and cannot be detected by click chemistry), and the benzyl alcohols **3a**-**3d** (which are released as part of the mechanism of action of the phosphate protecting group removal of **1a-1d** and **2a-2d**). EdU and an alkyne modified purine nucleoside analogue, EdA (7-deaza-7-ethynyl-2′-deoxyadenosine), were also included as positive controls. As expected (Ligasová et al., 2015), EdU did not inhibit HeLa cell growth at the highest concentration tested (100 µM) in either 4 h or 24 h assays (**Table 1**). EdA displayed low toxicity, with a 50% inhibitory concentration (IC_50_) of 54.38 (± 8.11) µM, in line with a previous report (Neef et al., 2012). Acyloxybenzyl acetyl analogue **1a** and the corresponding thymidine analogue **2a** displayed no toxicity with IC_50_ values >50 µM (**Table 1**). In contrast, the acyloxybenzyl benzoyl, pivaloyl and heptanoyl analogues (**1b-1d** and **2b-2d**) had IC_50_’s of 5.10-9.48 µM and 3.69-10.33 µM, respectively, in 4 h assays (**Table 1**) and 3.35-5.63 µM and 2.80-5.21 µM, respectively, in 24 h assays (**Table 1**). The reactive quinone methide released with loss of the acyloxy benzyl phosphate masking groups may lead to the higher relative toxicity (Meier et al., 2002) seen in the DNADetect™ probes (**1b-1d**) and control compounds (**2b-2d**) as compared to EdU. The corresponding benzyl alcohols **3a**-**3d** did not display toxicity in 4 h or 24 h assays (IC_50_>50 µM; **Table 1**). This was not surprising as these compounds are unlikely to generate quinone methide intermediates as the hydroxyl being a poor leaving group suppresses the umpolung mediated elimination reaction.

**Table 1.**
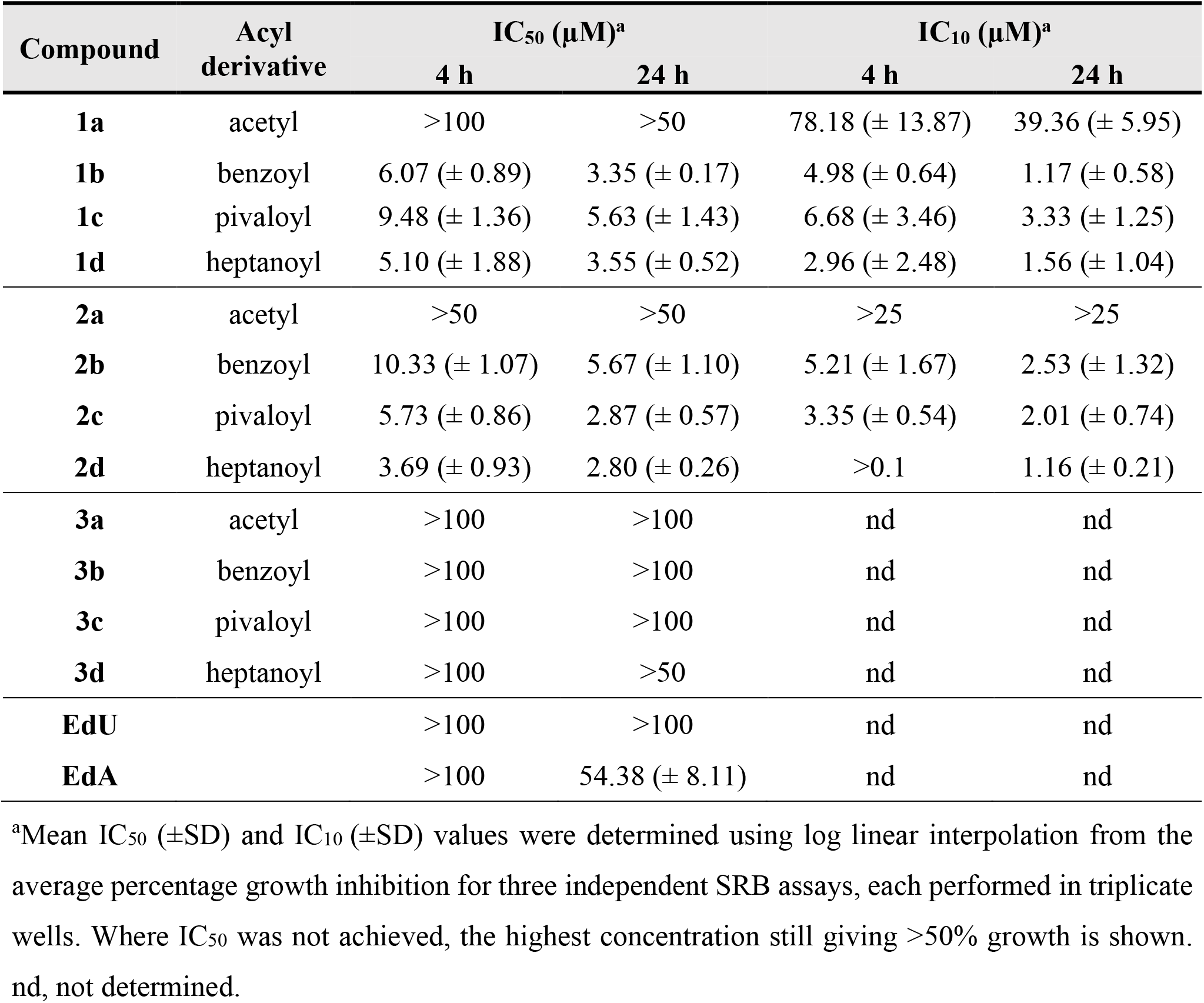
HeLa cell cytotoxicity of DNADetect™ probes 1a-d and corresponding chemical probe controls (2a-2d, 3a-3d).

| Compound | Acyl derivative | $IC_{50} (\mu M)^a$ | | $IC_{10} (\mu M)^a$ | |
| --- | --- | --- | --- | --- | --- |
|  |  | 4 h | 24 h | 4 h | 24 h |
| **1a** | acetyl | >100 | >50 | 78.18 ($\pm 13.87$) | 39.36 ($\pm 5.95$) |
| **1b** | benzoyl | 6.07 ($\pm 0.89$) | 3.35 ($\pm 0.17$) | 4.98 ($\pm 0.64$) | 1.17 ($\pm 0.58$) |
| **1c** | pivaloyl | 9.48 ($\pm 1.36$) | 5.63 ($\pm 1.43$) | 6.68 ($\pm 3.46$) | 3.33 ($\pm 1.25$) |
| **1d** | heptanoyl | 5.10 ($\pm 1.88$) | 3.55 ($\pm 0.52$) | 2.96 ($\pm 2.48$) | 1.56 ($\pm 1.04$) |
| **2a** | acetyl | >50 | >50 | >25 | >25 |
| **2b** | benzoyl | 10.33 ($\pm 1.07$) | 5.67 ($\pm 1.10$) | 5.21 ($\pm 1.67$) | 2.53 ($\pm 1.32$) |
| **2c** | pivaloyl | 5.73 ($\pm 0.86$) | 2.87 ($\pm 0.57$) | 3.35 ($\pm 0.54$) | 2.01 ($\pm 0.74$) |
| **2d** | heptanoyl | 3.69 ($\pm 0.93$) | 2.80 ($\pm 0.26$) | >0.1 | 1.16 ($\pm 0.21$) |
| **3a** | acetyl | >100 | >100 | nd | nd |
| **3b** | benzoyl | >100 | >100 | nd | nd |
| **3c** | pivaloyl | >100 | >100 | nd | nd |
| **3d** | heptanoyl | >100 | >50 | nd | nd |
| **EdU** |  | >100 | >100 | nd | nd |
| **EdA** | | >100 | 54.38 ($\pm 8.11$) | nd | nd |

<sup>a</sup>Mean $IC_{50}$ ( $\pm SD$ ) and $IC_{10}$ ( $\pm SD$ ) values were determined using log linear interpolation from the average percentage growth inhibition for three independent SRB assays, each performed in triplicate wells. Where $IC_{50}$ was not achieved, the highest concentration still giving >50% growth is shown. nd, not determined.

### Quantitative analysis of DNADetect™ probe labelling using flow cytometry

Flow cytometry was employed to quantify incorporation of **1a-1d** in comparison to EdU. HeLa cells were seeded at 40,000 cells/mL and 70,000 cells/mL overnight and then incubated with each probe for 4 h or 24 h. EdU and DMSO vehicle were included as controls. The IC_10_ of each probe (**Table 1**) was used to determine labelling concentrations that would be non-toxic. Probes **1b-1d** were tested at 1 µM. As **1a** gave the lowest toxicity, this probe was also tested at 5 µM. EdU was included at 1 µM and 5 µM. Following exposure to each probe or control, cells were harvested, fixed in 2% paraformaldehyde and 0.2% glutaraldehyde, and permeabilized with 0.1% TritonX-100, before undergoing a CuAAC reaction with Alexa Fluor 488 azide (ThermoFisher, USA) to enable flow cytometry detection at 488 nm. Cells were also treated with Hoechst 33342 to label DNA (ThermoFisher, USA). Three washes with 1% BSA/PBS were performed between each step.

Following 4 h treatment with 1 µM or 5 µM EdU, the percentage of Alexa Fluor 488-positive cells (compared to the DMSO control) was 34.7% and 52.0%, respectively, and following 24 h treatment 63.5% and 91.9%, respectively (**Figure 1**; **Supplemental Table S1**). These data were in line with previously published work with EdU (Ligasova et al., 2016). In comparison, 15.4%, 13.1%, 30.3% and 20.1% of the HeLa cells were Alexa Fluor 488-positive following 4 h labelling with 1 µM **1a-d**, respectively, and 38.5% of cells were labelled by 5 µM **1a** (**Figure 1**; **Supplemental Table S1**). This level of 4 h labelling was like that obtained with the cyclosal phosphotriester pronucleotide analogue of EdU previously developed by our group (Huynh et al., 2015), however in that case a significantly higher concentration of the probe was required (40 µM). As with EdU, higher labelling was obtained using probes **1a-d** in 24 h assays, with 75.5%, 66.3%, 75.2% and 58.4% of cells being Alexa Fluor 488 positive, respectively, and at 5 µM, 94.8% of cells were labelled by **1a** (**Figure 1**; **Supplemental Table S1**). As expected for the thymidine-equivalent controls **2a**-**2d**, which lack the alkyne group required for click chemistry detection, no Alexa Fluor 488-positive cells were detected after 24 h (<1% Alexa Fluor 488-positive cells compared to DMSO control, **Supplemental Table S1**).

**Figure 1.**
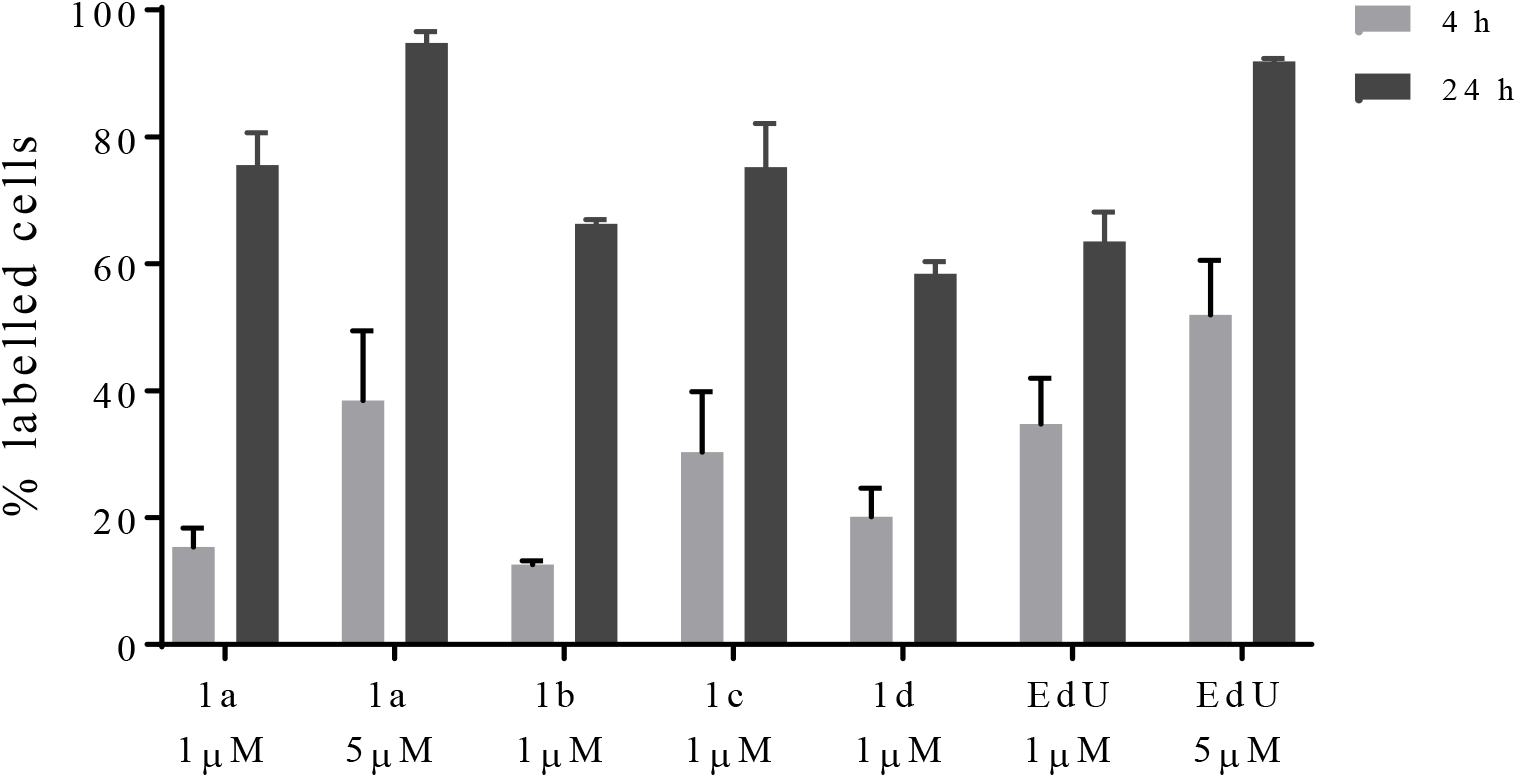
Flow cytometry analysis of DNADetect™ probes 1a-1d incorporation into HeLa cells. Mean (± SD) percentage of Alexa Fluor 488 and Hoechst 33342 co-labelled cells treated with **1a**-**1d** in 4 h and 24 h assays, compared to the positive control EdU and normalized to the DMSO control. At least two independent experiments (≥5,000 events counted per experiment) were performed.

### Visualization of DNADetect™ probes 1a-1d incorporation via fluorescence microscopy

To investigate the cellular location of DNADetect™ probes **1a-1d**, HeLa cells were seeded onto 12 mm glass coverslips in 24 well plates (Griener, USA) and exposed to 1 µM EdU or **1a-1d** for 24 h. Cells were fixed and permeabilized in-plate as per flow cytometry experiments, followed by CuAAC with Alexa Fluor 488 azide (on parafilm), and then returned to 24 well plates and stained with Hoechst 33342. Coverslips were then mounted onto microscope slides and assessed via fluorescence microscopy. Alexa Fluor 488 fluorescence was observed in cells treated with DNADetect™ probes **1a-1d** and EdU, but not in the negative control DMSO (**Figure 2**). Alexa Fluor 488 staining in **1a-1d** treated cells co-localized with Hoechst 33342 staining (**Figure 2**) indicated incorporation into DNA. The labelling intensity of each DNADetect™ probe was comparable to that displayed by EdU.

**Figure 2.**
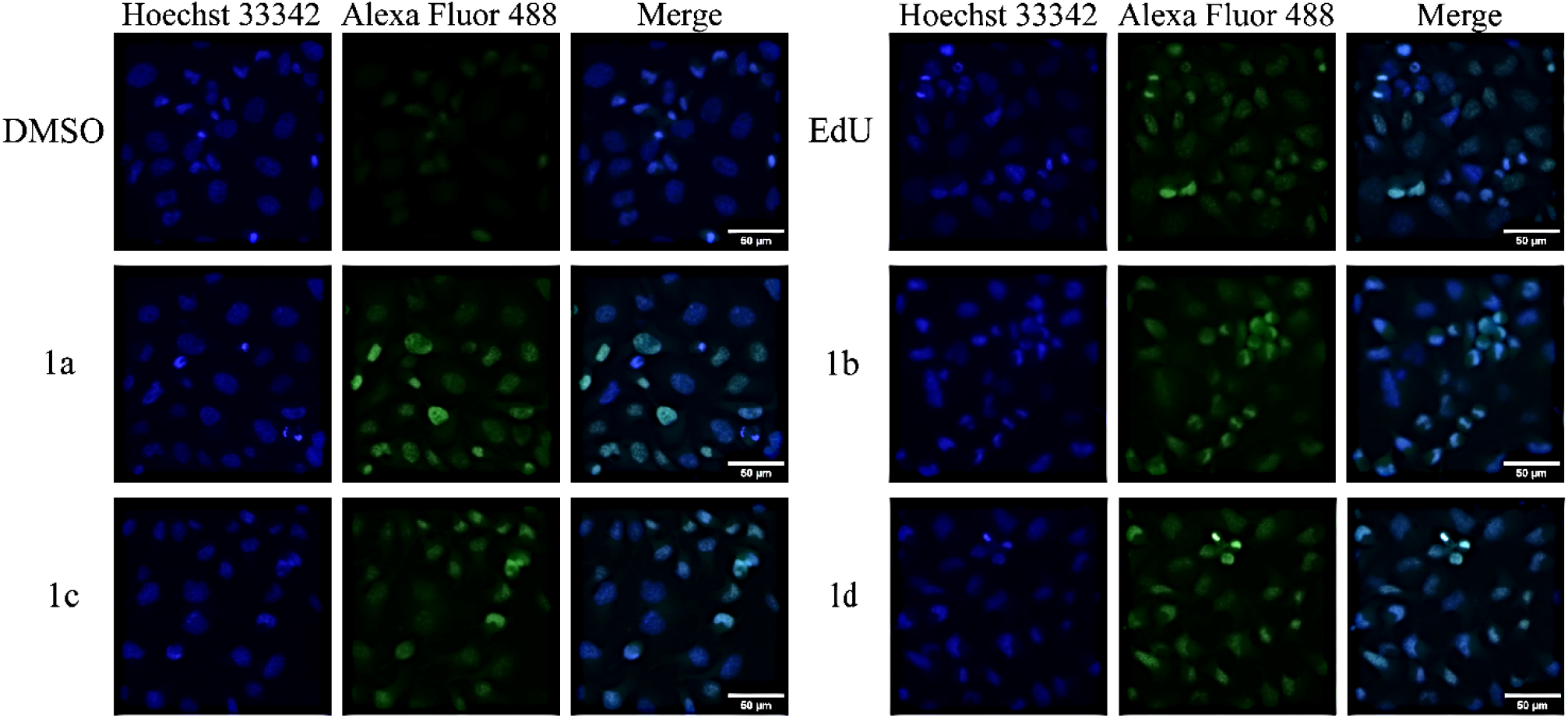
DNADetect™ probes 1a-1d label HeLa cell nuclei. Fluorescence microscopy of HeLa treated with DMSO vehicle control or 1 µM of **1a-1d** or EdU for 24 h and labelled with Alexa Fluor 488 azide (green) and Hoechst 33342 (blue). Merged images are shown. Images obtained on an Evident BX63 microscope with a 20⨯ objective.

### DNADetect™ probe 1d labelling is outcompeted by thymidine

EdU labelling efficiency has been shown to be reduced by thymidine in competition biolabeling studies (Ligasová et al., 2015), as both nucleosides are actively transported into the cell and undergo phosphorylation by thymidine kinase (Bitter et al., 2020; Wright & Lee, 2021). To assess the effect of thymidine competition on DNADetect™ probe labelling, HeLa cells were incubated for 8 h with 1 µM thymidine. Following CuAAC reactions with Alexa Fluor 488 azide and Hoechst 33342 DNA staining, cells were analysed using flow cytometry. Treatment with 2 µM EdU or **1d** for 8 h resulted in 54.7% and 53.3% of cells being detected as Alexa Fluor 488 positive, respectively (**Figure 3**; **Supplemental Table S2**). When cells were co-incubated with 1 µM thymidine, labelling was reduced by 80-90% for both EdU (consistent with a previous report (Ligasová et al., 2015)) and **1d** (**Figure 3**; **Supplemental Table S2**) in comparison to the DMSO control. These data demonstrate that thymidine can compete with both EdU and **1d** and impact labelling with these probes.

**Figure 3.**
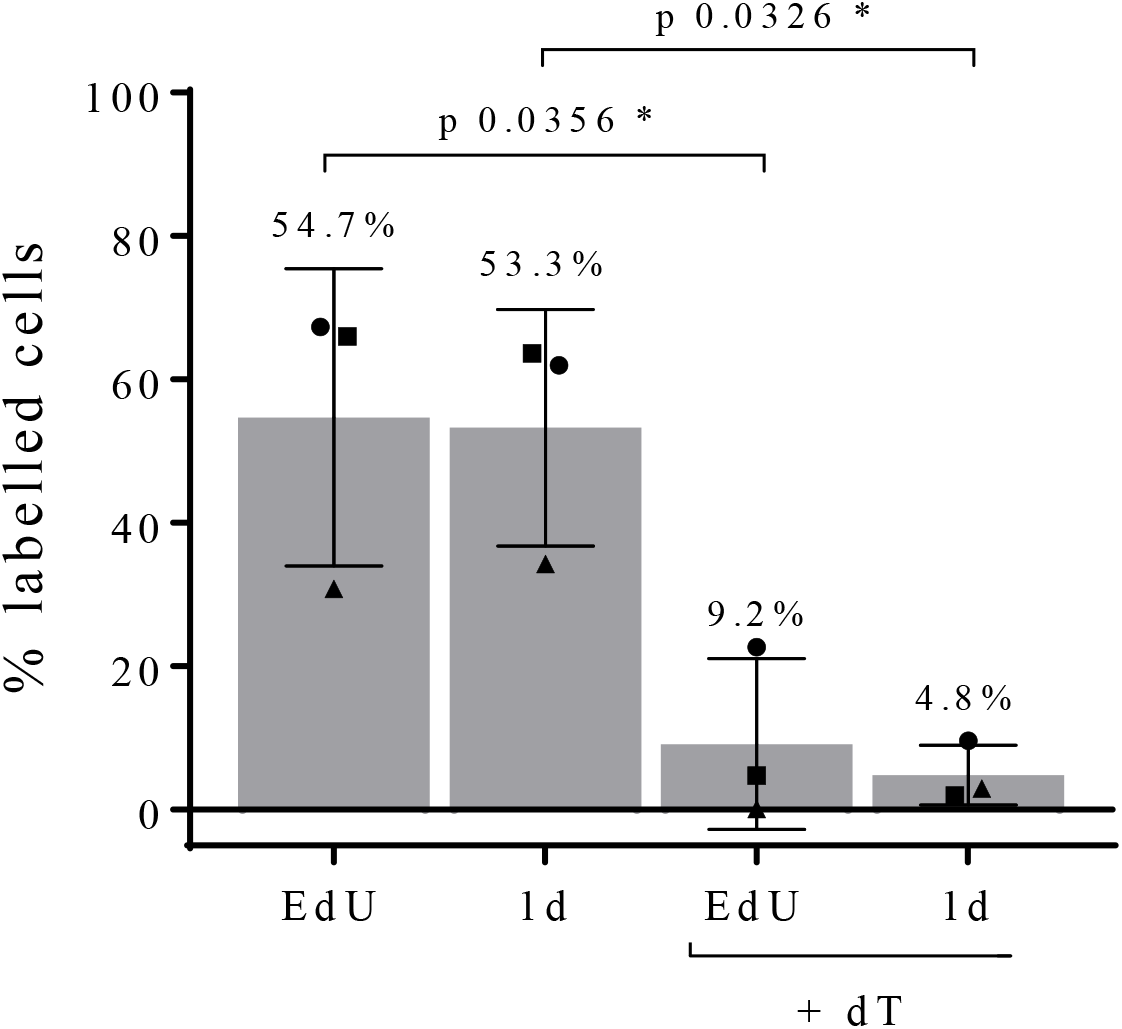
DNA labelling using EdU and DNADetect™ probe 1d is reduced in the presence of thymidine. Labelling efficiency was determined by flow cytometry of HeLa cells treated with 2 µM EdU or **1d** for 8 h in the presence of 1 µM thymidine (dT). Data are mean (± SD) percentage of Alexa Fluor 488 and Hoechst 33342 labelled cells from three independent experiments (≥5,000 events counted per experiment), normalized to respective DMSO and thymidine (dT) only controls (i.e., no EdU or **1d**).

## Conclusions

Nucleoside analogue probes are important biolabeling tools to detect DNA replication and synthesis (reviewed in (Solius et al., 2021)). In previous work, we designed EdU-based DNADetect™ probes **1b-1d** to passively diffuse through cellular membranes due to their lipophilicity and subsequently unmask intracellularly to reveal the monophosphate form of EdU (EdU-P_1_). We demonstrated these probes can be used to detect replicating DNA in *P. falciparum* malaria parasites which lack thymidine kinase by bypassing the requirement for monophosphorylation that is required for EdU incorporation (Hilko et al., 2023). Here, we have demonstrated that probes **1b-1d** can also label replicating DNA in HeLa cells. Furthermore, we have shown that like EdU, the incorporation of **1d** into HeLa cell DNA is inhibited by thymidine. Together these data demonstrate that DNADetect™ probes **1b-1d** are biolabeling tools that can be applied not only to applications where EdU is used but may also have broader applicability for thymidine transporter-independent and thymidine kinase-independent applications, such as when these may be inhibited or defective. Ideally, future work will evaluate the specific transport and incorporation mechanisms in these situations.

## Methods

### Mammalian cell culture and *in vitro* cytotoxicity

HeLa (Gey et al., 1954) were cultured in RPMI 1640 media (Gibco, USA) supplemented with penicillin-streptomycin 5% (Gibco, USA) and foetal bovine serum 10% (Bovagen, NZ), at 37 °C in 5% CO_2_. Harvesting cells to seed for use in cytotoxicity assays or labelling experiments consisted of a gentle rinse with warm PBS 1x (Sigma, USA) and disassociation (trypsin-EDTA 0.25%, Gibco, USA) for 2-3 min. Cells were collected in media, centrifuged at 300 G for 5 min, and resuspended in media for a cell count with trypan blue (Gibco, USA). Harvested HeLa were seeded into 96-well plates at a density of 7,000 cells/well and 4,000 cells/well for 4 h and 24 h cytotoxicity assays, respectively, overnight before the addition of DNADetect™ probes **1a-1d** and controls. Controls included EdU, EdA, chloroquine, thymidine-equivalent probes **2a-2d** and respective benzyl alcohol prodrug masking groups **3a-3d**. Following incubation at 37 °C, assay plates were gently emptied and rinsed once with warm PBS 1x (Sigma, USA). Cells were then fixed with methylated spirits (Recochem, Australia) for at least 30 min then rinsed once with RO water and allowed to airdry briefly. Sulforhodamine B (0.4% in 1% acetic acid, 50 µL) was added to each well and incubated at room temperature for at least 1 h, then rinsed three times with 1% acetic acid. Finally, 100µL tris buffer (10mM) was added to each well and incubated for at least 30 min before absorbance at 540 nm was read using a Biotek Synergy 2 (Agilent, USA). Percent inhibition was determined by comparing growth of treated cells to cell-only wells, minus background. IC_50_ values were determined in Excel using log-linear interpolation (Huber & Koella, 1993).

### Flow cytometry and fluorescence microscopy

HeLa were seeded in T75 flasks (Corning, USA) at 70,000 cells/mL or 40,000 cells/mL for 4 h and 24 h labeling experiments, respectively. Following an 8-12 h incubation, cultures were exposed to test and control compounds for 4 h or 24 h at 37 °C. A vehicle only negative control (≤0.0625% DMSO; non-toxic to the cells) was also included. For thymidine competition experiments, cells were seeded at 70,000 cells/mL in T25 flasks and incubated with 5 mL of 2 µM EdU, 2 µM probe 1d or equivalent vehicle for 8 h. Each flask was competed with either 1 µM thymidine or DMSO (final 0.02%, non-toxic).

Following treatment, cells were harvested as previously described, however were further washed once with cold PBS then immediately fixed in 2% paraformaldehyde (Sigma, USA) and 0.2% glutaraldehyde (Sigma, USA) for 15 min at room temperature. Cells were then washed twice by resuspending in 2 mL 1% bovine serum albumin (BSA; Bovagen, NZ) in PBS and centrifugation for 5 min at 300 G in an Eppendorf 5702 centrifuge (Eppendorf, Germany). The resulting pellet was permeabilized for 10 min at room temperature in 0.1% Triton^®^X-100 (Sigma, USA) in PBS. After washing twice with 1% BSA/PBS, as above, cell pellets were stained for 30 min at room temperature protected from light, using a CuAAC reaction cocktail (0.15-0.5 mL) containing 5 µM Alexa Fluor 488 azide (ThermoFisher, USA), 1 mM CuSO_4_(aq) (Sigma, USA) and 100 mM aqueous sodium ascorbate (Sigma, USA). The cells were then washed twice in 1% BSA/PBS and co-stained with 0.5mL 2 µM Hoechst 33342 (Thermo Fisher, USA) for 15 min at room temperature, protected from light. Following two final washes in 1% BSA/PBS, samples were resuspended in 0.3-0.5 mL 1% BSA/PBS before acquisition on a MACSQuant^®^ flow cytometer (Miltenyi, USA) were mounted on microscope slides with Prolong™ Glass Antifade (ThermoFisher Scientific, USA). Alexa Fluor 488 fluorescence was measured using a blue laser at 488 nm and band pass filter of 525/50 nm. Hoechst 33342 was measured using a violet laser at 405 nm and band pass filter 450/50 nm. The voltage settings and acquisition parameters were as follows, Forward Scatter (FSC) 498v, linear mode; Side Scatter (SSC) 548v, linear mode; Alexa Fluor 488 and Hoechst 33342 400v, hLog. Doublets were excluded by gating on FSC-Area (FSC-A) and FSC-Height (FSC-H) and 5,000 events were counted per sample. Data were analyzed with FlowJo (version 10, Becton, Dickinson & Company, USA) and reported as the mean percentage ±SD (n ≥ 2) of Alexa Fluor 488/Hoechst 33342 positive cells normalized to the DMSO negative control.

For fluorescence microscopy, images were acquired using an Evident BX63 fluorescence microscope (Evident, USA) with cellSens software (Olympus, USA) using a 20’ objective. Excitation and emission wavelengths used were Hoechst 33342 Ex: 325-375, Em: 435-485; Alexa Fluor 488 Ex: 450-490, Em: 500-550. Exposure times to acquire all images were Hoechst 33342 (5.415 ms) and Alexa Fluor 488 (1.415 ms). Images were exported into FIJI software (version 1.53t) for cropping and scale bar labelling.

## Supporting Information

Description of synthetic methods and full characterization including ^1^H, ^13^C, ^31^P NMR spectra for all compounds.

## Acknowledgments

This research was supported by the Australian Government through the Australian Research Council’s Discovery Projects funding scheme (projects DP220102618, DP180102601).

## Notes

### Competing Interest Statement

The authors have declared no competing interest.

## References

Bitter, E. E., Townsend, M. H., Erickson, R., Allen, C., & O’Neill, K. L. (2020). Thymidine kinase 1 through the ages: a comprehensive review. Cell Biosci, 10(1), 138. 10.1186/s13578-020-00493-1

Buck, S. B., Bradford, J., Gee, K. R., Agnew, B. J., Clarke, S. T., & Salic, A. (2008). Detection of S-phase cell cycle progression using 5-ethynyl-2’-deoxyuridine incorporation with click chemistry, an alternative to using 5-bromo-2’-deoxyuridine antibodies. Biotechniques, 44(7), 927–929. 10.2144/000112812

Djordjevic, B., & Szybalski, W. (1960). Genetics of human cell lines. III. Incorporation of 5-bromo- and 5-iododeoxyuridine into the deoxyribonucleic acid of human cells and its effect on radiation sensitivity. J Exp Med, 112(3), 509–531. 10.1084/jem.112.3.509

Eidinoff, M. L., Cheong, L., & Rich, M. A. (1959). Incorporation of unnatural pyrimidine bases into deoxyribonucleic acid of mammalian cells. Science, 129(3362), 1550–1551. 10.1126/science.129.3362.1550

Farre, X., Guillen-Gomez, E., Sanchez, L., Hardisson, D., Plaza, Y., Lloberas, J., Casado, F. J., Palacios, J., & Pastor-Anglada, M. (2004). Expression of the nucleoside-derived drug transporters hCNT1, hENT1 and hENT2 in gynecologic tumors. Int J Cancer, 112(6), 959–966. 10.1002/ijc.20524

Gey, G. O., Bang, F. B., & Gey, M. K. (1954). Responses of a variety of normal and malignant cells to continuous cultivation, and some practical applications of these responses to problems in the biology of disease. Annals of the New York Academy of Sciences, 58(7), 976–999. 10.1111/j.1749-6632.1954.tb45886.x

Gratzner, H. G. (1982). Monoclonal antibody to 5-bromo- and 5-iododeoxyuridine: A new reagent for detection of DNA replication. Science, 218(4571), 474–475. 10.1126/science.7123245

Hilko, D. H., Fisher, G. M., Addison, R. S., Andrews, K. T., & Poulsen, S. A. (2023). Thymidine Kinase-Independent Click Chemistry DNADetect Probes for DNA Proliferation Assessment in Malaria Parasites. ACS Chem Biol, 18(12), 2535–2543. 10.1021/acschembio.3c00530

Huber, W., & Koella, J. C. (1993). A comparison of three methods of estimating EC50 in studies of drug resistance of malaria parasites. Acta Trop, 55(4), 257–261. 10.1016/0001-706x(93)90083-n

Huynh, N., Dickson, C., Zencak, D., Hilko, D. H., Mackay-Sim, A., & Poulsen, S. A. (2015). Labeling of Cellular DNA with a Cyclosal Phosphotriester Pronucleotide Analog of 5-ethynyl-2’-deoxyuridine. Chem Biol Drug Des, 86(4), 400–409. 10.1111/cbdd.12506

Ligasova, A., & Koberna, K. (2018). DNA Replication: From Radioisotopes to Click Chemistry. Molecules, 23(11). 10.3390/molecules23113007

Ligasova, A., Liboska, R., Friedecky, D., Micova, K., Adam, T., Ozdian, T., Rosenberg, I., & Koberna, K. (2016). Dr Jekyll and Mr Hyde: a strange case of 5-ethynyl-2’-deoxyuridine and 5-ethynyl-2’-deoxycytidine. Open Biol, 6(1), 150172. 10.1098/rsob.150172

Ligasová, A., Strunin, D., Friedecky, D., Adam, T., & Koberna, K. (2015). A Fatal Combination: A Thymidylate Synthase Inhibitor with DNA Damaging Activity. Plos One, 10(2). 10.1371/journal.pone.0117459

Marti-Clua, J. (2024). 5-Bromo-2’-deoxyuridine labeling: historical perspectives, factors influencing the detection, toxicity, and its implications in the neurogenesis. Neural Regen Res, 19(2), 302–308. 10.4103/1673-5374.379038

Meier, C., Muus, U., Renze, J., Naesens, L., De Clercq, E., & Balzarini, J. (2002). Comparative Study of Bis(Benzyl)Phosphate Triesters of 2′,3′-Dideoxy-2′,3′-Didehydrothymidine (d4T) and CycloSal-d4TMP — Hydrolysis, Mechanistic Insights and Anti-HIV Activity. Antiviral Chemistry and Chemotherapy, 13(2), 101–114. 10.1177/095632020201300204

Neef, A. B., Samain, F., & Luedtke, N. W. (2012). Metabolic labeling of DNA by purine analogues in vivo. Chembiochem, 13(12), 1750–1753. 10.1002/cbic.201200253

Pastor-Anglada, M., & Perez-Torras, S. (2018). Emerging Roles of Nucleoside Transporters. Front Pharmacol, 9, 606. 10.3389/fphar.2018.00606

Perez-Torras, S., Iglesias, I., Llopis, M., Lozano, J. J., Antolin, M., Guarner, F., & Pastor-Anglada, M. (2016). Transportome Profiling Identifies Profound Alterations in Crohn’s Disease Partially Restored by Commensal Bacteria. J Crohns Colitis, 10(7), 850–859. 10.1093/ecco-jcc/jjw042

Perez-Torras, S., Vidal-Pla, A., Cano-Soldado, P., Huber-Ruano, I., Mazo, A., & Pastor-Anglada, M. (2013). Concentrative nucleoside transporter 1 (hCNT1) promotes phenotypic changes relevant to tumor biology in a translocation-independent manner. Cell Death Dis, 4(5), e648. 10.1038/cddis.2013.173

Pradere, U., Garnier-Amblard, E. C., Coats, S. J., Amblard, F., & Schinazi, R. F. (2014). Synthesis of nucleoside phosphate and phosphonate prodrugs. Chem Rev, 114(18), 9154–9218. 10.1021/cr5002035

Sawicki, D. L., Erlanger, B. F., & Beiser, S. M. (1971). Immunochemical detection of minor bases in nucleic acids. Science, 174(4004), 70–72. 10.1126/science.174.4004.70

Skehan, P., Storeng, R., Scudiero, D., Monks, A., McMahon, J., Vistica, D., Warren, J. T., Bokesch, H., Kenney, S., & Boyd, M. R. (1990). New colorimetric cytotoxicity assay for anticancer-drug screening. J Natl Cancer Inst, 82(13), 1107–1112. 10.1093/jnci/82.13.1107

Solius, G. M., Maltsev, D. I., Belousov, V. V., & Podgorny, O. V. (2021). Recent advances in nucleotide analogue-based techniques for tracking dividing stem cells: An overview. J Biol Chem, 297(5), 101345. 10.1016/j.jbc.2021.101345

Wright, N. J., & Lee, S. Y. (2021). Toward a Molecular Basis of Cellular Nucleoside Transport in Humans. Chem Rev, 121(9), 5336–5358. 10.1021/acs.chemrev.0c00644

Xu, X., Li, Z., Yao, X., Sun, N., & Chang, J. (2023). Advanced prodrug strategies in nucleoside analogues targeting the treatment of gastrointestinal malignancies. Front Cell Dev Biol, 11, 1173432. 10.3389/fcell.2023.1173432

Young, J. D., Yao, S. Y., Baldwin, J. M., Cass, C. E., & Baldwin, S. A. (2013). The human concentrative and equilibrative nucleoside transporter families, SLC28 and SLC29. Mol Aspects Med, 34(2-3), 529–547. 10.1016/j.mam.2012.05.007

