## Supplemental Information for "Thymidine kinase-independent click chemistry DNADetect™ probes as an EdU alternative for mammalian cell DNA labelling"

**Supplementary Information**

**Table S1. Flow cytometry analysis of HeLa cells labelled with DNADetect™ probes**

| Compound | Alexa Fluor 488 Positive cells | |
|---|---|---|
| | 4 h | 24 h |
| **1a 1 µM** | 15.4% (± 2.9) | 75.5% (± 5.1) |
| **1a 5 µM** | 38.5% (± 11.0) | 94.8% (± 1.8) |
| **1b 1 µM** | 13.1% (± 1.0) | 66.3% (± 0.7) |
| **1c 1 µM** | 30.3% (± 9.5) | 75.2% (± 6.9) |
| **1d 1 µM** | 20.2% (± 4.5) | 58.4% (± 1.9) |
| **2a 1 µM** | nd | 0.3% (± 0.8) |
| **2a 5 µM** | nd | -0.1% (± 0.4) |
| **2b 1 µM** | nd | -0.1% (± 0.2) |
| **2c 1 µM** | nd | 0.1% (± 0.02) |
| **2d 1 µM** | nd | -0.1% (± 0.4) |
| **EdU 1 µM** | 34.7% (± 7.3) | 63.5% (± 4.7) |
| **EdU 5 µM** | 52.0% (± 8.6) | 91.9% (± 0.4) |

The labelling efficiency of each probe over 4 h and 24 h was normalized to the DMSO control and is presented as mean (± SD) percentage of Alexa Fluor 488 and Hoechst 33342 labelled cells from at least two independent experiments (≥5,000 events counted per experiment). nd, not determined.

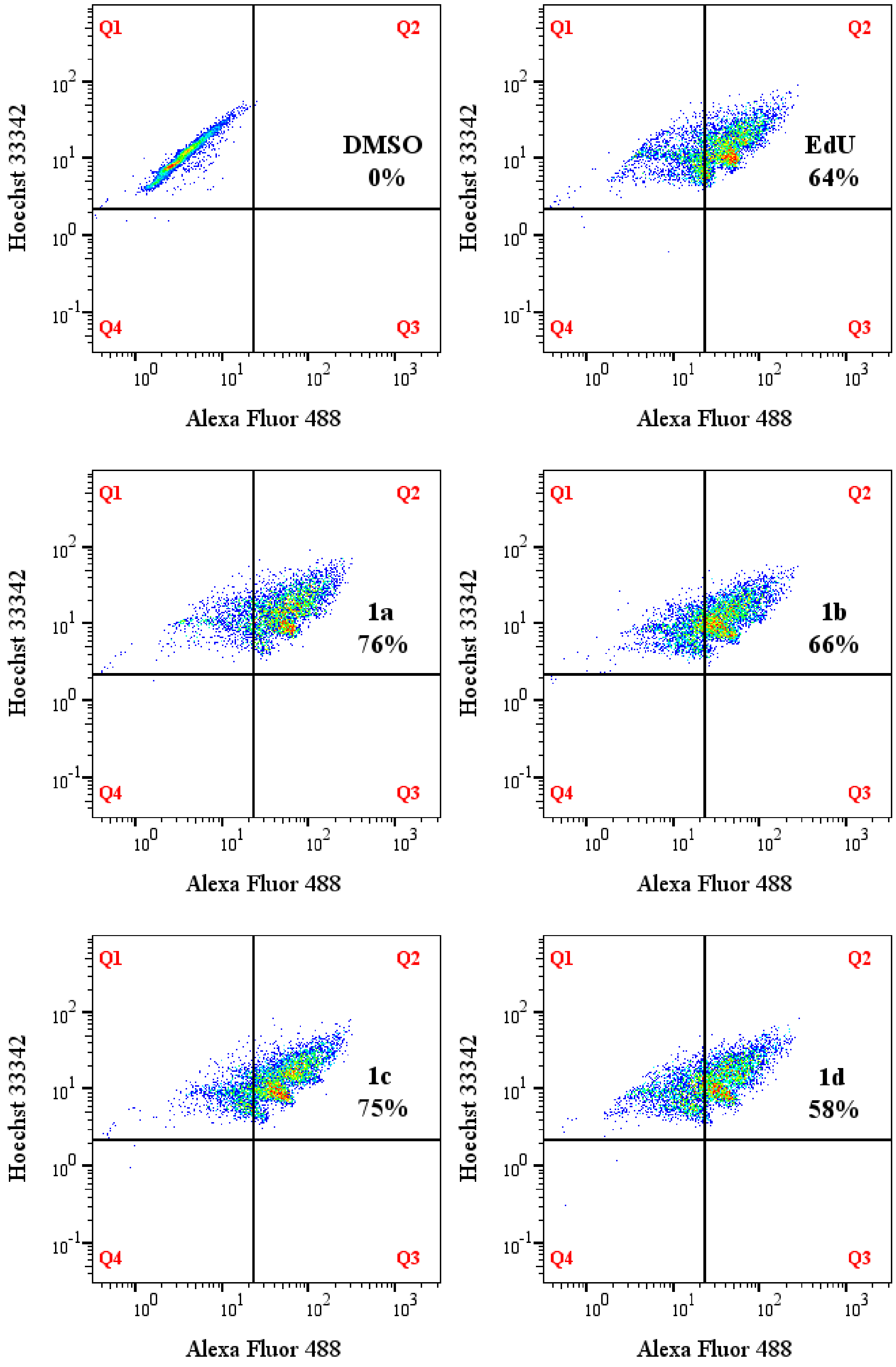

**Figure S1. Labelling efficiency of DNADetect probes in HeLa.** Representative flow cytometry dot plots one independent experiment showing Hoechst 33342 and Alexa Fluor 488 staining of HeLa following 24 h of exposure to 1 µM DNADetect probes **1a–1d,** EdU and DMSO vehicle control. Population gates are denoted as Q1, Hoechst 33342 only; Q2, Hoechst 33342 + Alexa Fluor 488; Q3, Alexa Fluor 488 only; and Q4, unstained. Percentages annotated in gate Q2 indicate Hoechst 33342- and Alexa Fluor 488-positive cells.

**Table S2. Probe 1d incorporation is diminished in the presence of thymidine.**

| Probe | Alexa Fluor 488 Positive cells | |
|---|---|---|
| | **(-)** | **(+) 1 μM dT** |
| **EdU** | 54.7% (± 20.7) | 9.2% (± 11.9) |
| **1d** | 53.3% (± 16.5) | 4.8% (± 4.2) |

Labelling efficiency of each probe (2 μM) following 8 h exposure is presented as mean (± SD) percentage of Alexa Fluor 488 and Hoechst 33342 labelled cells normalized to DMSO control data. At least two independent experiments (≥5,000 events counted per experiment) were performed.

#### General Chemistry

All reactions were carried out in dry solvents under anhydrous conditions unless otherwise stated. All chemicals were purchased from commercial suppliers and used without further purification. All reactions were monitored by TLC using silica plates with visualisation of eluted bands by UV fluorescence (λ = 254 nm) and charring with vanillin stain (6 g vanillin in 100 mL of EtOH containing 1% v/v 98% Sulfuric acid. Silica gel flash chromatography was performed using silica gel 60 Å (230-400 mesh). NMR ($^1$H, $^{13}$C, COSY, NOESY, HSQC and HMBC) spectra were recorded on either a Bruker AVANCE III HD 500 MHz NMR spectrometer equipped with a BBO probe at 25 °C or a Bruker AVANCE III HD 400 MHz NMR spectrometer equipped with a BBO probe at 25 °C. Chemical Shifts for $^1$H and $^{13}$C NMR obtained in DMSO-$d_6$ are reported in ppm relative to residual solvent proton (d = 2.50 ppm) and carbon (d = 39.5 ppm) signals, respectively. Chemical Shifts for $^1$H and $^{13}$C NMR obtained in Methanol-$d_4$ are reported in ppm relative to residual solvent proton (d = 3.31 ppm) and carbon (d = 49.0 ppm) signals, respectively. Signal splitting multiplicity is indicated as follows: s (singlet), d (doublet), t (triplet), q (quartet), m (multiplet), dd (doublet of doublets), br (broad signal), a (apparent). Assignments of $^1$H and $^{13}$C chemical shifts were established by COSY, HSQC, HMBC, and NOESY experiments. Coupling constants are reported in hertz (Hz). LRMS (ESI) data were acquired on a Thermo Fisher MSQ Plus single quadruple ESI mass spectrometers using electrospray as the ionisation technique in positive and/or negative mode as stated. HRMS (ESI) data were acquired on a Bruker MaXis II QTOF mass spectrometer using an ESI source the ionisation technique in positive-ion and/or negative mode as stated. All MS analysis samples were prepared as solutions in either methanol or acetonitrile. Purity of the compounds were >95% as determined by Thermo Fisher Dionex Ultimate 3000 series HPLC via UV detection at 254 nm. EdU, compounds **1a-1d** and phosphoramidites were synthesised as previously described (Hilko et al., 2023).

#### General Procedure

Thymidine was dissolved in pyridine then cooled to approximately -10 °C and then TFA added. The mixture was allowed to warm to RT then 3 Å molecular sieves were added. The mixture was sealed and stored at RT for at least 20 h. The reaction vessel (4 mL vial) was dried via heat gun (~300 °C) then flushed with argon. Next, the thymidine mixture was transferred to the reaction vessel and the mixture cooled to -20 °C. The phosphoramidite was dissolved in dry pyridine (10 equiv.), cooled to -20 °C then added dropwise to the reaction mixture with vigorous stirring. The reaction mixture was stirred at -20 °C for 1 h and allowed to warm to RT and stirred for an additional 1 hour. The reaction mixture was then cooled to -5 °C then *t*-BuOOH (5-6 M in decanes, 2.4 equiv) was added, allowed to warm to RT and stirred for 30 min. The reaction mixture was concentrated *in vacuo* and the residue dissolved in 10% MeOH/$CH_2Cl_2$ (10 mL) then stirred with a mixed bed ion exchange resin until universal pH paper indicated neutral pH and pyridine was undetectable by TLC (5% MeOH in $CH_2CH_2$) (~ 30 min). The mixed bed resin comprised Amberlyst® A-21 resin (20-30 mesh) and Dowex® 50WX8 resin ($H^+$-form. 100-200 mesh), both prewashed with MeOH, dried, then mixed via vortexing. The mixture was then filtered, washed with MeOH (15 mL), and concentrated *in vacuo*. The crude compound was purified by silica gel flash chromatography employing eluent mixtures and gradients as described for each compound.

**Compound 2a: Bis(4-acetyloxybenzyl) thymidine monophosphate**

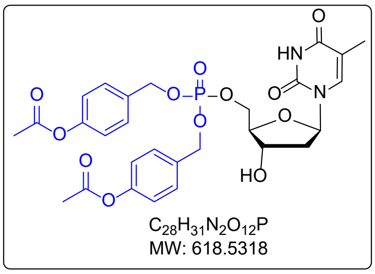

Following the general procedure, treatment of thymidine (1.0 equiv, 0.100 g, 0.4128 mmol) in pyridine (30 equiv, 1.385 mmol, 1.002 mL) with TFA (4.0 equiv, 1.651 mmol, 0.1264 mL), *N,N*-diisopropyl bis (4-acetyloxybenzyl)phosphoramidite (1.1 equiv, 0.266 g, 0.454 mmol) and *t*-BuOOH (5-6 M in decane, 2.4 equiv, 0.1801 mL), followed by silica gel flash column chromatography afforded the final product **2a** as a white solid (0.1007 g, 36%). $R_f$ (5% MeOH/ $CH_2Cl_2$) = 0.41. **$^1$H NMR (400 MHz, DMSO-*d*$_6$)** δ = 11.30 (br. s, 1H, N*H*-3), 7.46 (d, *J* = 1.3 Hz, 1H, H-6), 7.42 – 7.36 (m, 4H, 4× H-Bn$_{ortho}$), 7.18 – 7.06 (m, 4H, 4 x H-Bn$_{meta}$), 6.19 (a t, *J* = 6.9 Hz, 1H, H-1′), 5.43 (d, *J* = 4.3 Hz, 1H, OH-3′), 5.04 (d, $^3J$ = 8.0 Hz, 4H, 2× BnC$H_2$), 4.26 – 4.20 (m, 1H, H-3′), 4.20 – 4.07 (m, 2H, H-5′, H-5″), 3.94 - 3.88 (m, 1H, H-4′), 2.27 (s, 6H, 2×AcO), 2.12 – 2.04 (m, 2H, H-2′$_{(\alpha \text{ and } \beta)}$), 1.71(d, *J* = 1.2 Hz, 3H, C$H_3$-5). **$^{13}$C{H} NMR (101 MHz, DMSO-*d*$_6$)** δ = 169.16 (2× BnCO$_2$-), 163.66 (C-4), 150.49(2× C-Bn$_{para}$), 150.42 (C-2), 135.81 (C-6), 133.44 (d, $^3J_{C-P}$ = 6.9 Hz, 2× C-Bn$_{ipso}$), 129.19 (4×C-Bn$_{ortho}$), 121.94 (4× C-Bn$_{meta}$), 109.83 (C-5), 84.28 ($^3J_{C-P}$ = 7.3 Hz, C-4′), 83.88 (C-1′), 69.98 (C-3′), 68.12 (d, $^2J_{C-P}$ = 5.1 Hz, 2× Bn*C*H$_2$), 67.17(d, $^2J_{C-P}$ = 5.5 Hz, C-5′), 38.60 (C-2′), 20.85 (2× CO$_2$*C*H$_3$), 12.03 (*C*H$_3$-5). **$^{31}$P NMR (162 MHz, DMSO-*d*$_6$)** δ = -0.68 - -1.21 (m). **$^{31}$P{H} NMR (162 MHz, DMSO-*d*$_6$)** δ = -0.92 (s). ***m/z* (LRMS ESI$^+$)** 619.2 $[M + H]^+$. ***m/z* (LRMS ESI$^-$)** 617.4 $[M - H]^-$. ***m/z* (HRMS ESI$^-$) [M-H]$^-$** found 617.1542; calcd for $C_{28}H_{30}N_2O_{12}P$ 617.1537.

**Compound 2b: Bis(4-benzyloxybenzyl) thymidine monophosphate**

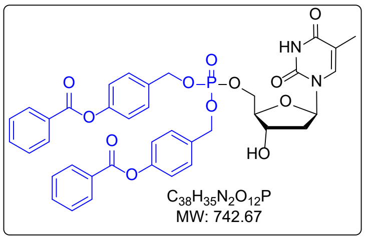

Following the general procedure, treatment of thymidine (1.0 equiv, 0.050 g, 0.2064 mmol) in pyridine (30 equiv, 6.192 mmol, 0.518 mL) with TFA (4.0 equiv, 0.8256 mmol, 0.063 mL), *N,N*-diisopropyl bis (4-(hydroxymethyl)phenyl benzoate)phosphoramidite (1.1 equiv, 0.133 g, 0.2270 mmol) and *t*-BuOOH (5-6 M in decane, 2.0 equiv, 0.0825 mL), followed by silica gel flash column chromatography afforded the final product **2b** as a white solid (0.035 g, 22.8%). $R_f$ (7.5% MeOH/ $CH_2Cl_2$) = 0.64. **$^1$H NMR (500 MHz, DMSO-$d_6$)** δ = 11.34 (s, br, 1H, NH), 8.17 – 8.11 (m, 4H, 4× H-Bz$_{ortho}$), 7.79 – 7.72 (m, 2H, 2× H-Bz$_{para}$), 7.65 – 7.57 (m, 4H, 4× H-Bz$_{meta}$), 7.50 (d, $J$ = 1.4 Hz, 1H, H-6), 7.50 – 7.47 (m, 4H, 4× H-Bn$_{ortho}$), 7.36 – 7.28 (m, 4H, 4× H-Bn$_{meta}$), 6.22 (a t, $J$ = 7.0 Hz, 1H, H-1′), 5.47 (d, $^3J$ = 4.2 Hz, 1H, OH-3′), 5.12 (m, 4H, 2× Bn$CH_2$), 4.30 – 4.16 (m, 3H, H-3′, H-5′, H-5″), 3.96 (dtd, $J$ = 5.0, 3.5, 1.2 Hz, 1H, H-4′), 2.17 – 2.07 (m, 2H, H-2′, H-2″), 1.74 (d, $^4J$ = 1.2 Hz, 3H, $CH_3$). **$^{13}$C{H} NMR (126 MHz, DMSO-$d_6$)** δ = 164.6 (2× *C*$O_2$Bn), 163.7 (C-4), 150.6 (2× Bn*C*O), 150.5 (C-2), 135.9 (C-6), 134.1 (2× C-Bz$_{para}$), 133.8 (d, $^3J_{CP}$ = 7.0 Hz, 2× C-Bn$_{ipso}$), 129.8 (4× C-Bz$_{meta}$), 129.3 (4× C-Bn$_{ortho}$), 129.0 (4× C-Bz$_{ortho}$), 128.8 (2× C-Bz$_{ipso}$), 122.1 (4× C-Bn$_{meta}$), 109.9 (C-5), 84.3 (d, $^3J$ = 7.3 Hz, C1′), 83.9 (C-4′), 70.0 (C-3′), 68.2 (d, $^3J_{CP}$ = 5.2 Hz, 2× Bn$CH_2$ ), 67.2 (d, $^3J$ = 5.7 Hz, C-5′), 38.6 (C-2′), 12.1 ($CH_3$). **$^{31}$P NMR (202 MHz, DMSO-$d_6$)** δ = -0.87 (p, $^3J$ = 7.4 Hz). **$^{31}$P{H} NMR (202 MHz, DMSO-$d_6$)** δ = -0.86 (s). ***m/z* (LRMS ESI$^+$)** 743.3 $[M + H]^+$, 765.3 $[M + Na]^+$. ***m/z* (LRMS ESI$^-$)** 741.2 $[M - H]^-$. ***m/z* (HRMS ESI$^+$) $[M+Na]^+$** found 765.1812; calcd **$[M+Na]^+$** for $C_{38}H_{35}N_2NaO_{12}P$ 765.1820.

**Compound 2c: Bis(4-*tert*butanoyloxybenzyl) thymidine monophosphate**

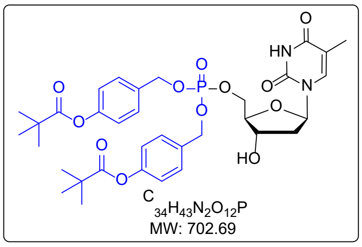

Following the general procedure, treatment of thymidine (1.0 equiv, 0.050 g, 0.2064 mmol) in pyridine (30 equiv, 6.192 mmol, 0.500 mL) with TFA (4.0 equiv, 0.8256 mmol, 0.063 mL), *N,N*-diisopropyl (Bis(4-tertbutyloxybenzyl)-*N,N*-diisopropylphosphoramidite) (1.1 equiv, 0.123 g, 0.2270 mmol) and *t*-BuOOH (5-6 M in decane, 2.0 equiv, 0.0825 mL), followed by silica gel flash column chromatography afforded the final product **2c** as a white solid (0.044 g, 32.0 %). $R_f$ (10% MeOH/ $CH_2Cl_2$) = 0.45.**$^1$H NMR (500 MHz, DMSO-$d_6$)** δ 11.33 (s, br, 1H, NH), 7.48 (d, $J$ = 1.2 Hz, 1H, H-6), 7.46 – 7.36 (m, 4H, 4x H-Bn$_{ortho}$), 7.14 – 7.06 (m, 4H, 4× H-Bn$_{\mathrm{meta}}$), 6.24 –

6.15 (m, 1H, H-1′), 5.46 (d, $^3J$ = 4.3 Hz, 1H, OH-3′), 5.06 (d, $J$ = 8.1 Hz, 4H, 2× Bn$CH_2$), 4.29 – 4.09 (m, 3H, H-3′, H-5′, H-5″), 3.93 (ddt, $J$ = 7.0, 5.0, 2.4 Hz, 1H, H-4′), 2.16 – 2.06 (m, 2H, 2′- H, H2″- H), 1.73 (d, $J$ = 1.2 Hz, 3H, $CH_3$), 1.30 (s, 18H, $(CH_3)_3$). **$^{13}$C{H} NMR (126 MHz, DMSO-*d$_6$*)** δ 176.4 (2× *C*$O_2$Bn), 163.7 (C-4), 150.8 (2× Bn*C*O), 150.4 (C-2), 135.8 (C-6), 133.4 (d, $^3J_{CP}$ = 6.7 Hz, 2× C-Bn$_{ipso}$), 129.3 (4× C-Bn$_{ortho}$), 121.8 (4× C-Bn$_{meta}$), 109.8 (C-5), 84.3 (d, $^3J$ = 7.3 Hz, C-1′), 83.9 (C-4′), 70.0 (C-3′), 68.2 (d, $^3J_{CP}$ = 5.3 Hz, 2× Bn$CH_2$), 67.2 (C-5′), 38.6 (*C*$(CH_3)_3$), 38.5 (d, $J$ = 5.6 Hz, C-2′), 26.8 (C(*C*$H_3)_3$), 12.1 (*C*$H_3$). **$^{31}$P NMR (202 MHz, DMSO-*d$_6$*)** δ -0.90 (p, $^3J$ = 30 Hz). **$^{31}$P NMR{H}** (202 MHz, DMSO-*d$_6$*) δ -0.88 (s). ***m/z* (LRMS ESI$^+$)** 703.3 $[M + H]^+$. ***m/z* (LRMS ESI$^-$)** 701.3 $[M - H]^-$. ***m/z* (HRMS ESI$^+$) [M+Na]$^+$** found 725.2452; calcd **[M+Na]$^+$** for $C_{34}H_{43}N_2NaO_{12}P$ 725.2470.

**Compound 2d: Bis(4-heptanoyloxybenzyl) thymidine monophosphate**

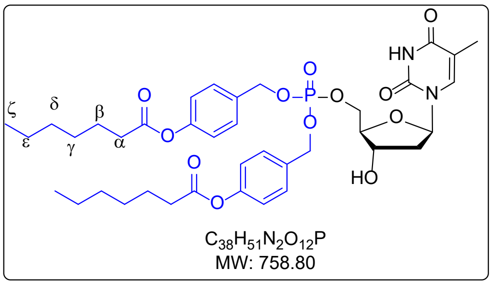

Following the general procedure, treatment of **thymidine**(1.0 equiv, 0.050 g, 0.2064 mmol) in pyridine (30 equiv, 6.192 mmol, 0.518 mL) with TFA (4.0 equiv, 0.8256 mmol, 0.063 mL), *N,N*-diisopropyl bis (4-(hydroxymethyl)phenyl heptanoate)phosphoramidite (1.1 equiv, 0.136 g, 0.2270 mmol) and *t*-BuOOH (5-6 M in decane, 2.0 equiv, 0.155 mL), followed by silica gel flash column chromatography afforded the final product **2d** as a white solid (0.032 g, 20.5%). $R_f$ (5% MeOH/ $CH_2Cl_2$) = 0.7. **$^1$H NMR (400 MHz, DMSO-*d$_6$*)** δ = 11.32 (s, br, 1H, NH), 7.48 (d, $^4J$ = 1.3 Hz, 1H, H-6), 7.44 – 7.37 (m, 4H, 2× H-Bn$_{ortho}$), 7.15 – 7.08 (m, 4H, 2× H-Bn$_{meta}$), 6.21 (a t, $^3J$ = 7.0 Hz, 1H, H-1′), 5.45 (s 1H, OH-3′), 5.05 (d, $J$ = 8.0 Hz, 4H, 2× Bn$CH_2$), 4.30 – 4.16 (m, 3H, H-3′, H-5′, H-5″), 3.93 (qd, $J$ = 4.0, 2.1 Hz, 1H, H-4′), 2.58 (t, $J$ = 7.4 Hz, 4H, 2× H-α-heptanoyl), 2.15-2.05 (m, 2H, 2′-H, 2″-H), 1.72 (d, $^4J$ = 1.3 Hz, 3H, $CH_3$), 1.64 (p, $J$ = 7.3, 7.2 Hz, 4H, 2× H-ß-heptanoyl), 1.38 – 1.28 (m, 12H, 2× H-γ-heptanoyl, H-δ-heptanoyl, H-ε-heptanoyl), 0.91 – 0.85 (m, 6H, 2× H-ζ-heptanoyl). **$^{13}$C NMR (101 MHz, DMSO-*d$_6$*)** δ 171.7 (2× *C*$O_2$Bn), 163.6 (C-4), 150.4 (d, $^3J$ = 6.5 Hz, 2× Bn*C*O), 135.8 (C-2), 133.4 (d, $^3J$ = 6.9 Hz, 2× C-Bn$_{ipso}$), 129.2 (4× C-Bn$_{ortho}$), 121.9 (4× C-Bn$_{meta}$), 109.8 (C-5), 84.3 (d, $J$ = 7.5 Hz, C4′), 83.9 (C-1′), 70.0 (C-3′), 68.1 (d, $^3J$ = 5.3 Hz, 2× Bn*C*$H_2$), 67.2 (C-5′), 38.6 (C-2′), 33.4 (C-α-heptanoyl), 30.9 (C-δ-heptanoyl), 28.1 (C-γ-heptanoyl), 24.3 (C-ß-heptanoyl), 21.9 (C-ε-heptanoyl), 13.9 (C-ζ-heptanoyl), 12.0 (*C*$H_3$). **$^{31}$P NMR (162 MHz, DMSO-*d$_6$*)** δ -0.91 (p, $^3J$ = 7.6Hz). **$^{31}$P{H} NMR (162 MHz, DMSO-*d$_6$*)** δ -0.91 (s). ***m/z* (LRMS ESI$^+$)** 759.5 [M

+ H $^+$. ***m/z* (LRMS ESI$^-$)** 757.5 [M - H] $^-$. ***m/z* (HRMS ESI$^+$) [M + Na]$^+$** found 781.3070; calcd **[M+Na]$^+$** for $C_{38}H_{51}N_2NaO_{12}P$ 781.3072.

### NMR for Compound 2a: Bis(4-acetyloxybenzyl) thymidine monophosphate

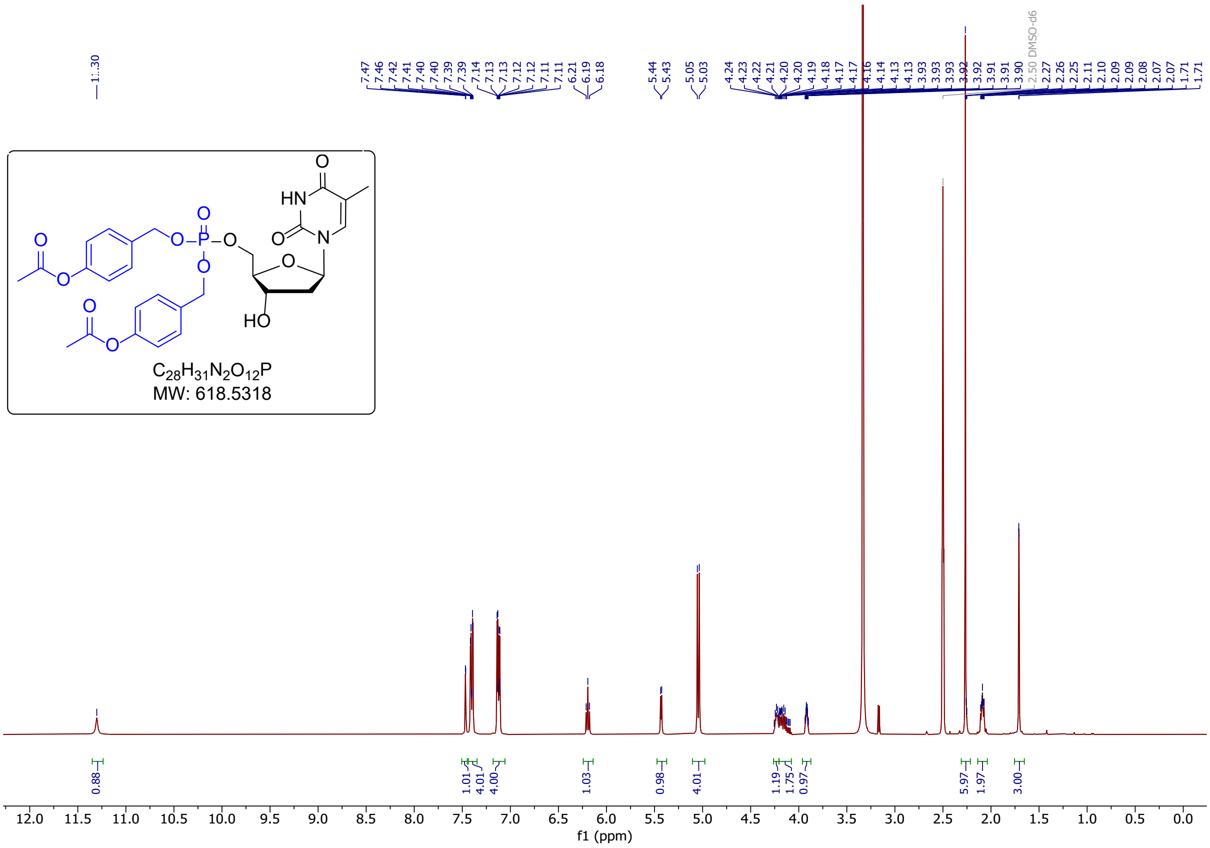

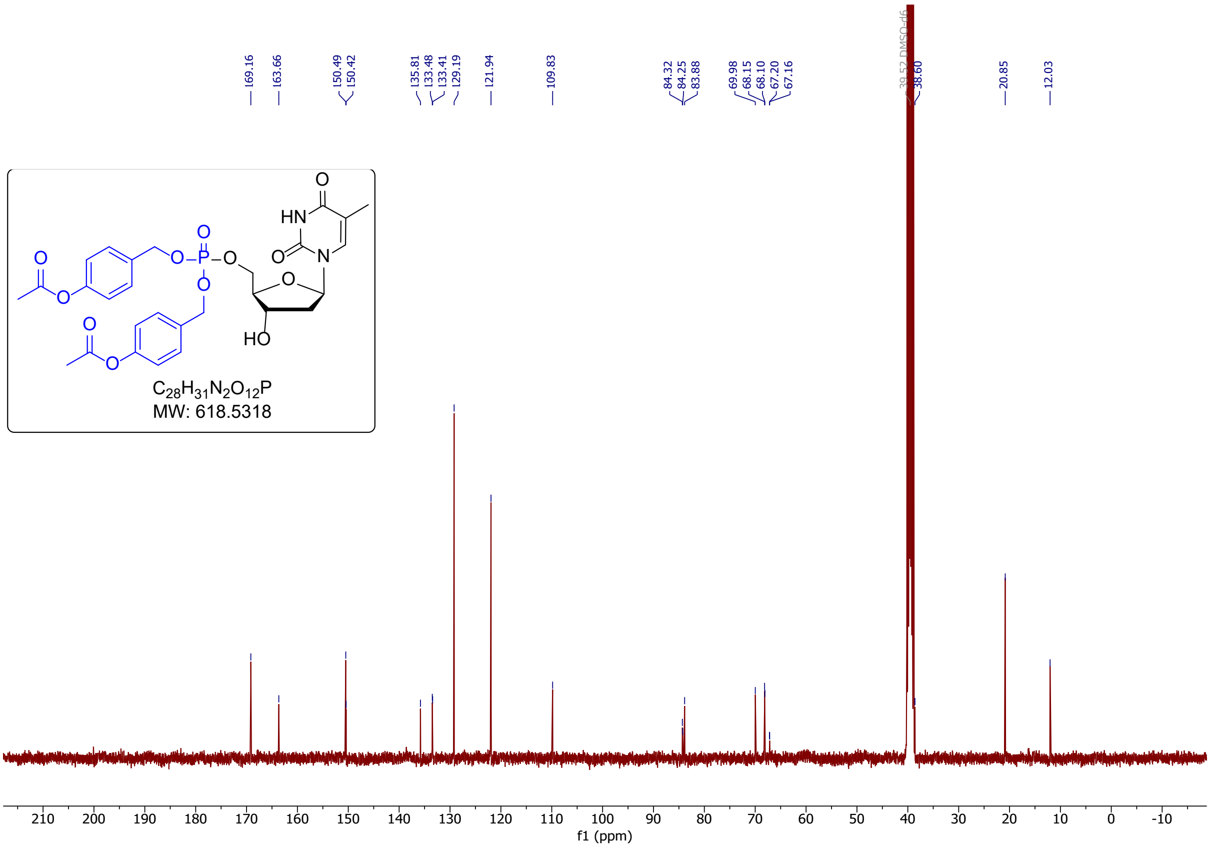

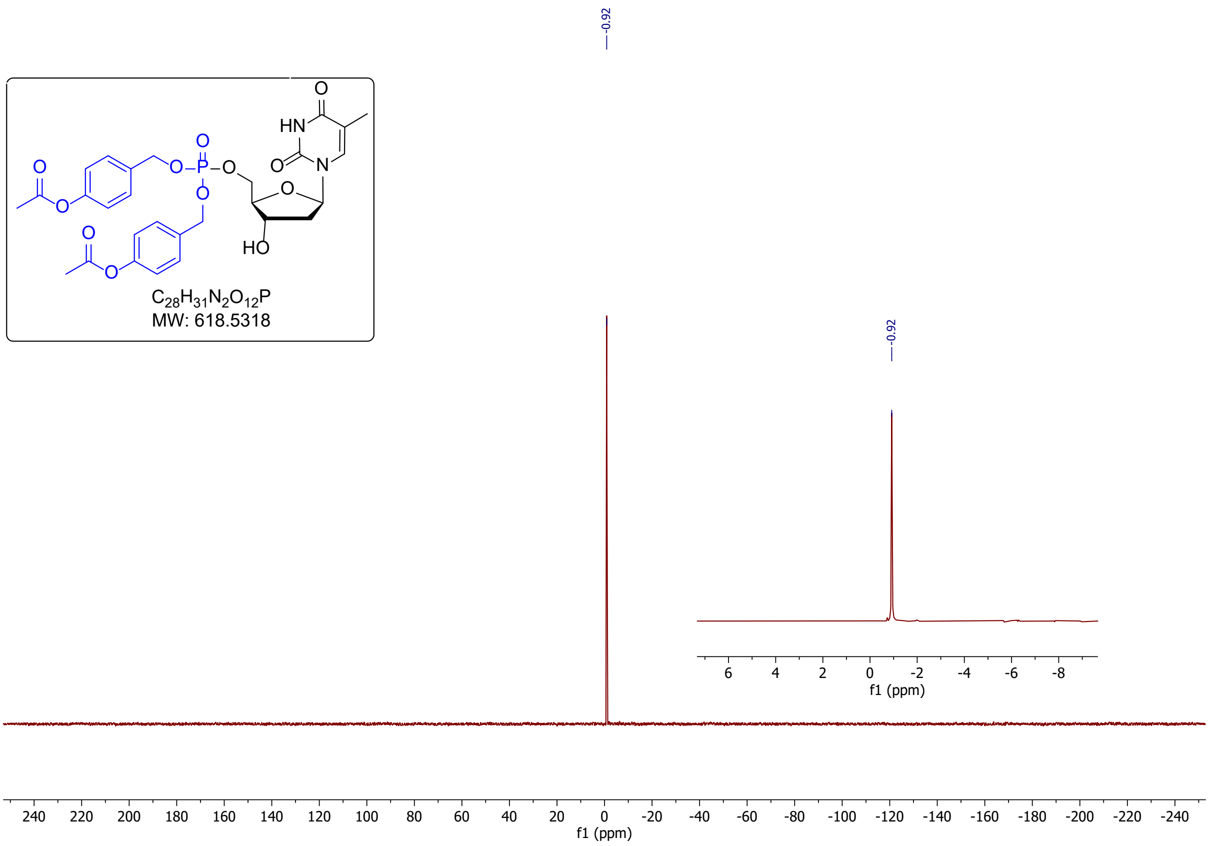

$^{31}P\{H\}$NMR (DMSO-$d_6$)

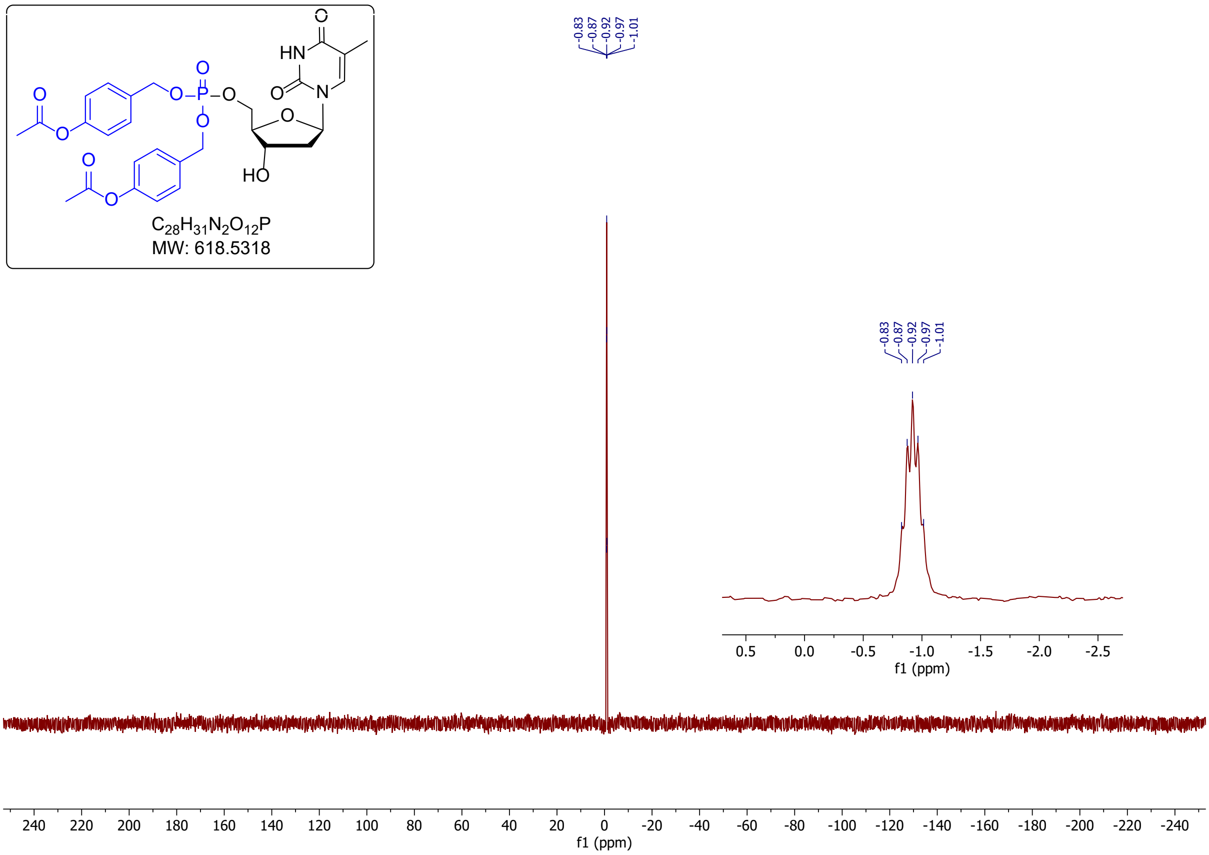

### NMR Spectra for compound 2b: bis(4-benzyloxybenzyl) thymidine monophosphate

$^{1}$HNMR (DMSO-$d_6$)

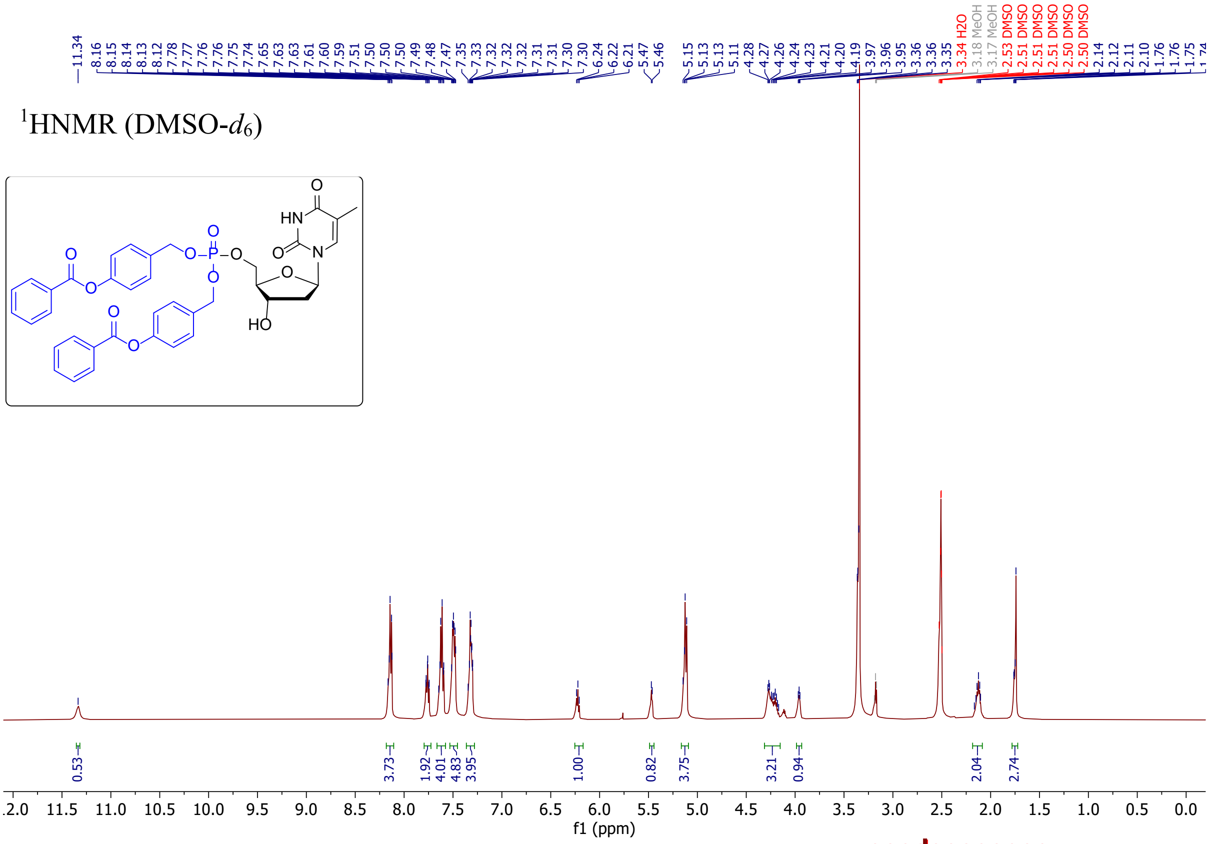

$^{13}$CNMR (DMSO-$d_6$)

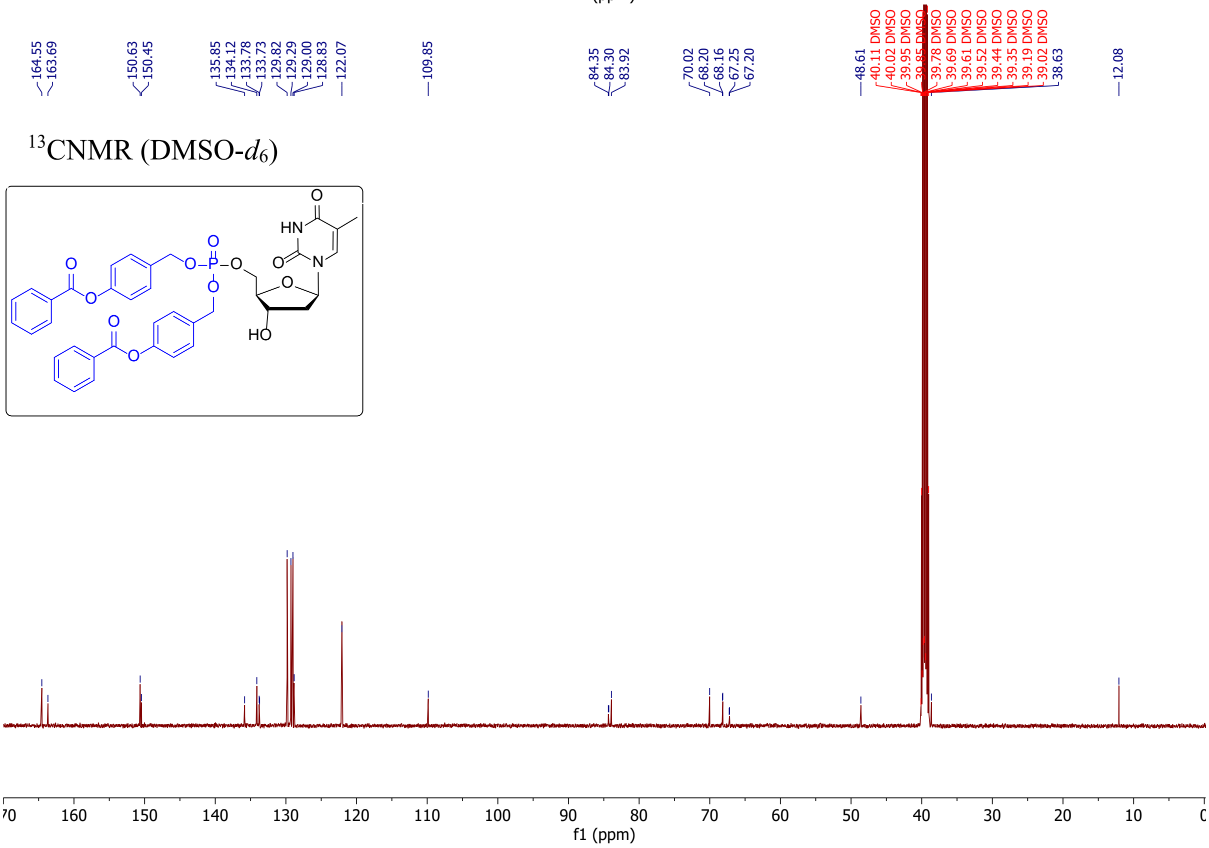

$^{31}$PNMR (DMSO-$d_6$)

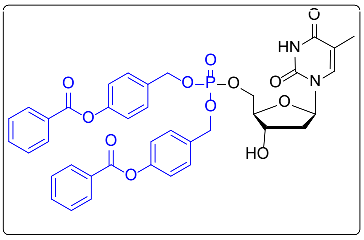

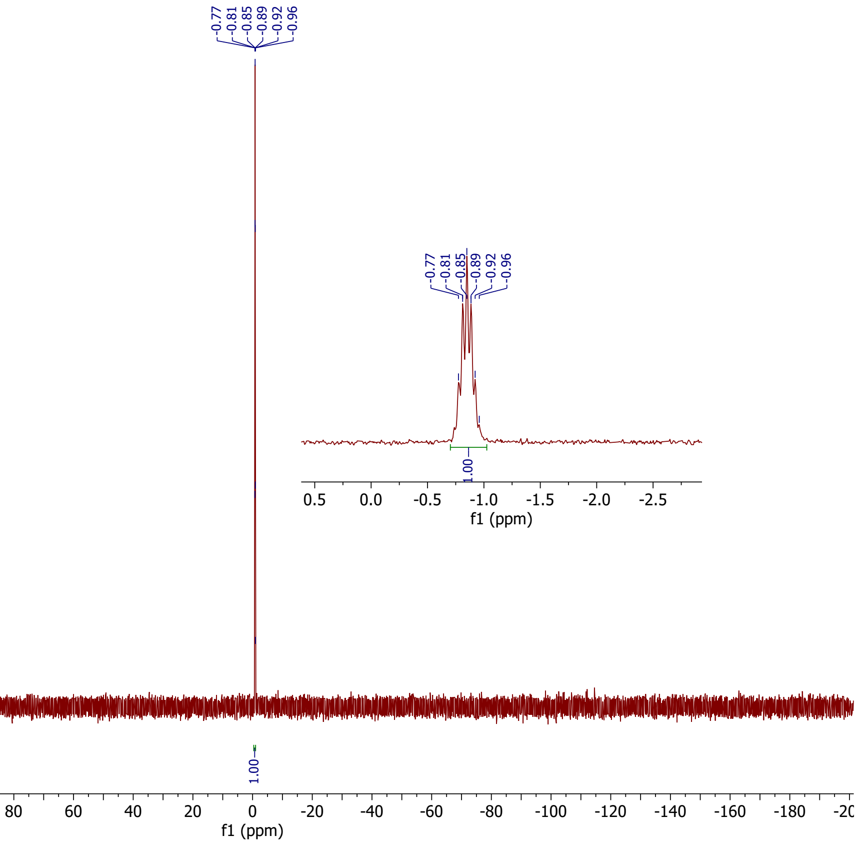

$^{31}$P{H}NMR (DMSO-$d_6$)

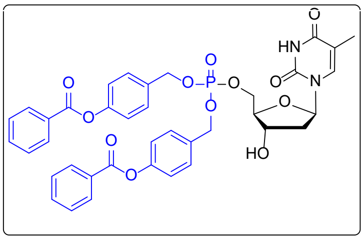

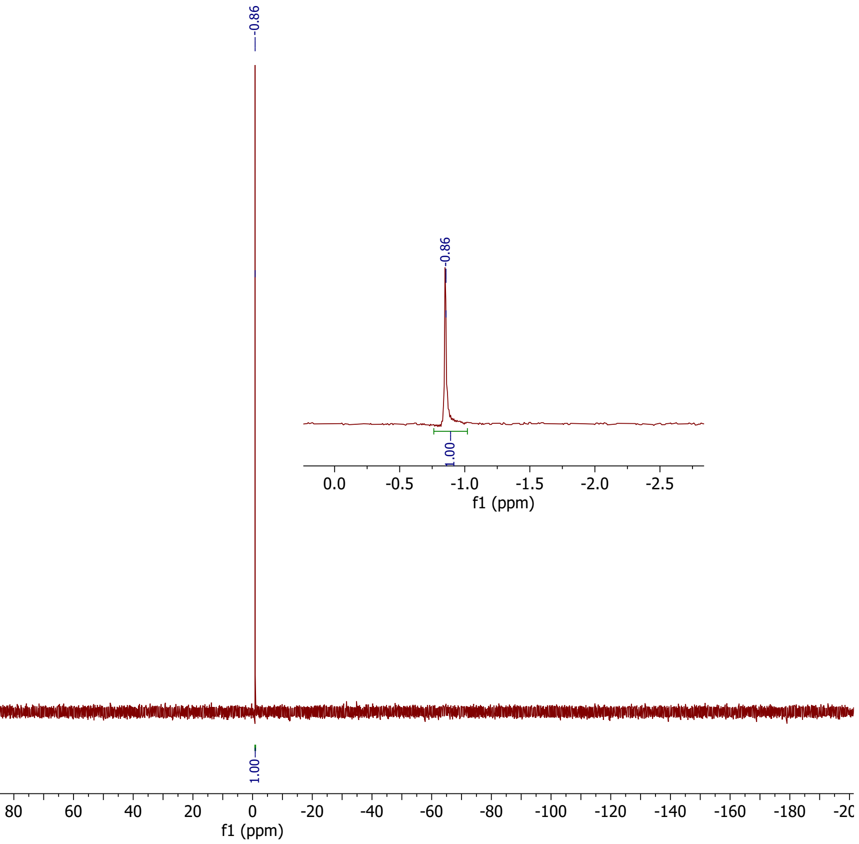

### NMR spectra for compound 2c: Bis(4-*tert*butanoyloxybenzyl)-thymidine monophosphate

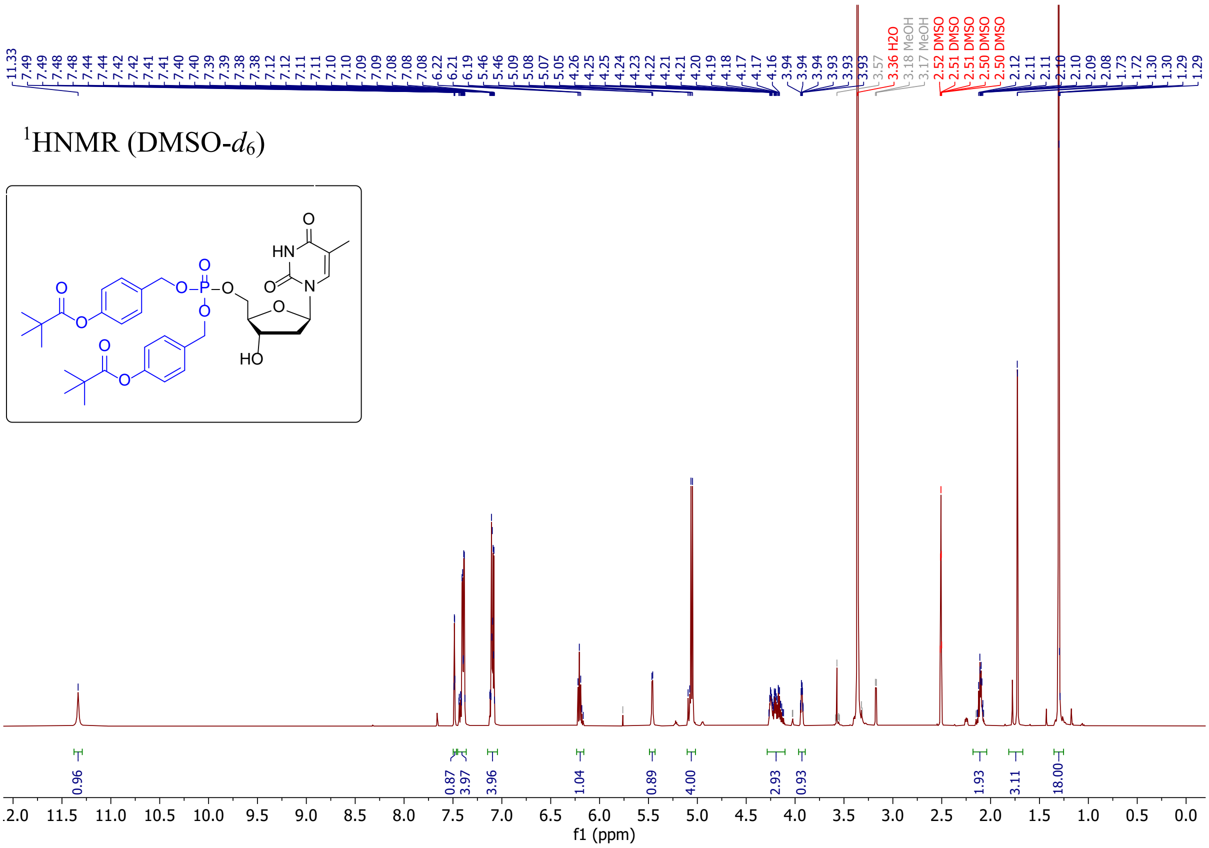

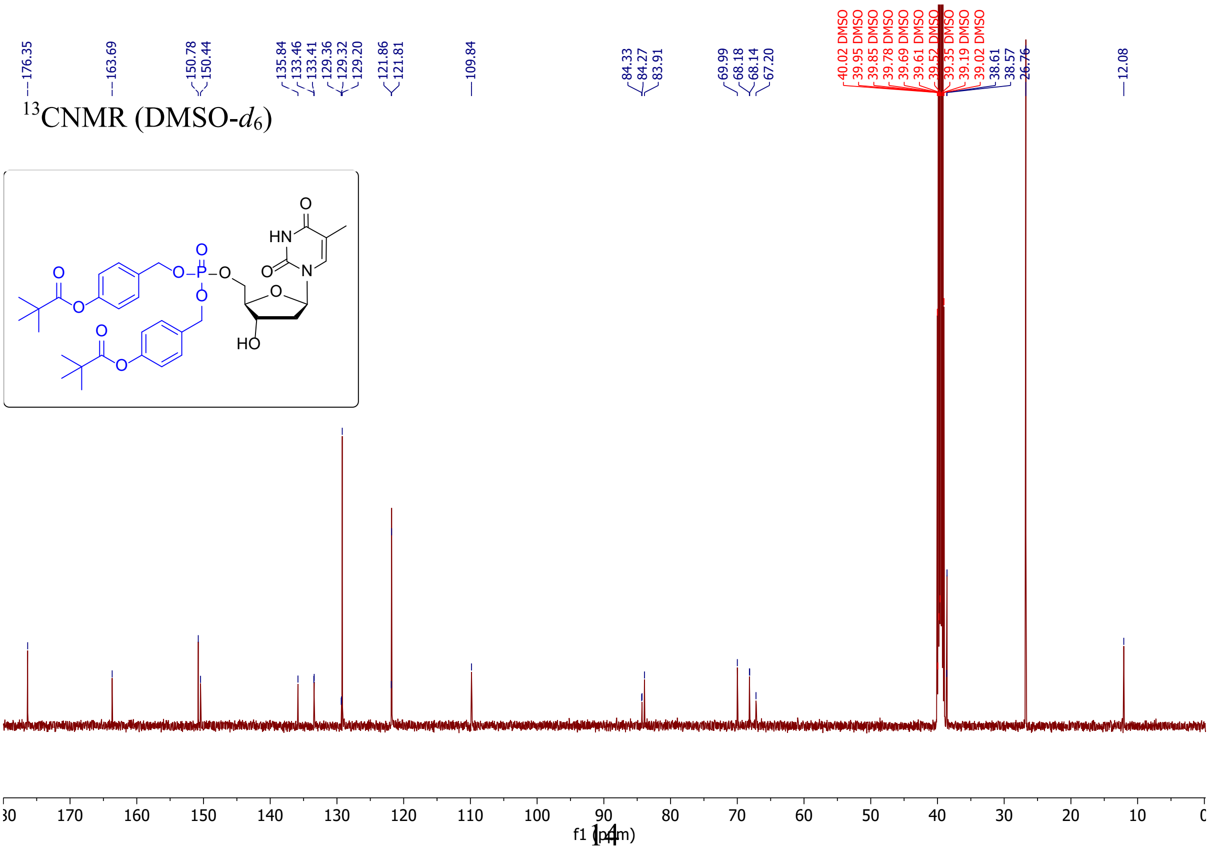

$^{31}$PNMR (DMSO-$d_6$)

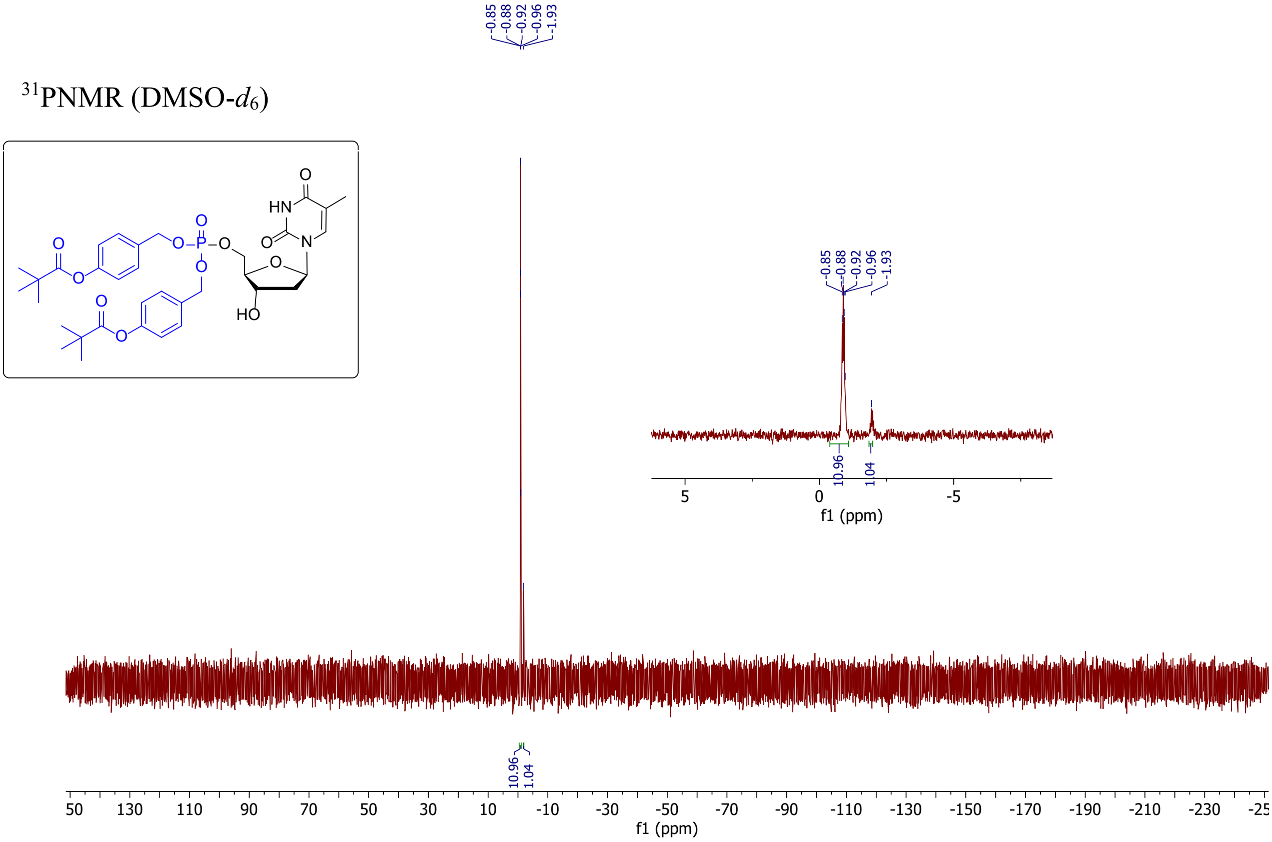

$^{31}$P{H}NMR (DMSO-$d_6$)

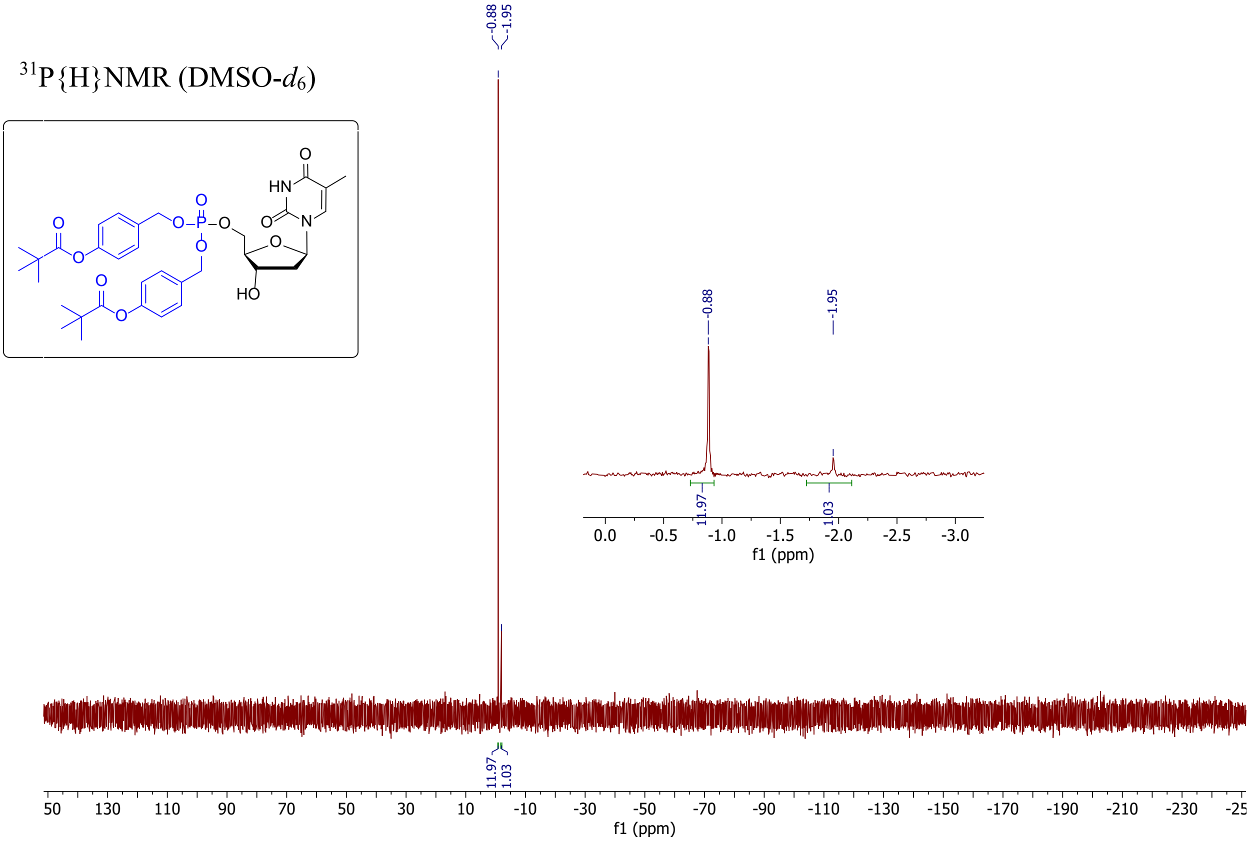

#### NMR spectra for compound 2d: Bis(4-heptanoyloxybenzyl) thymidine monophosphate

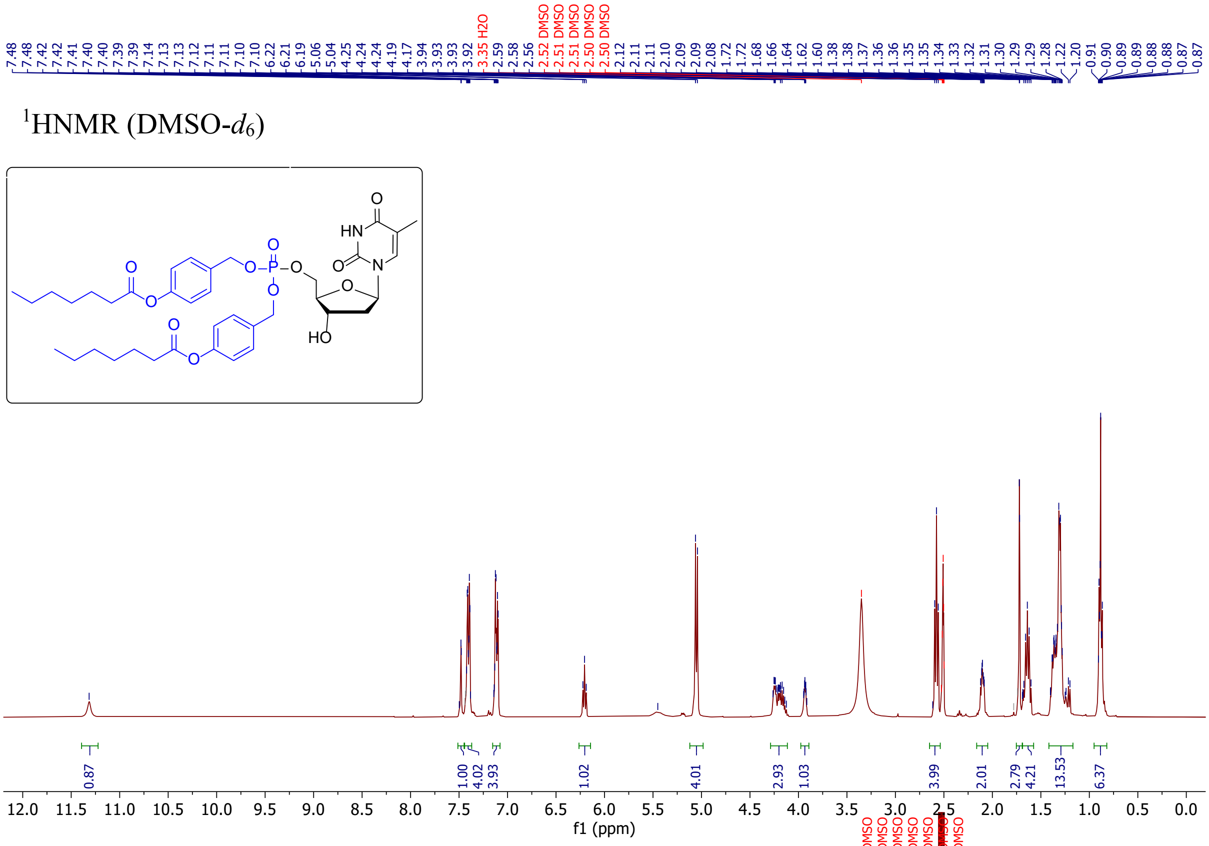

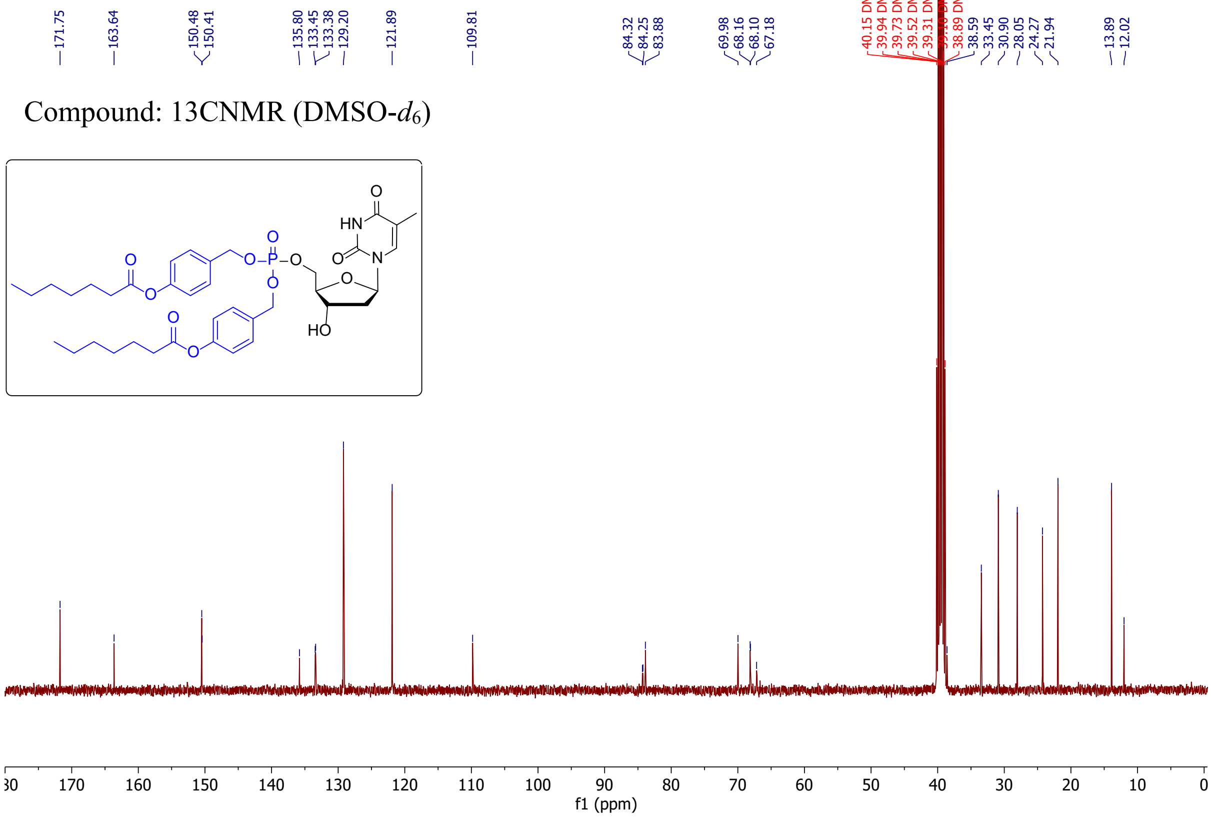

$^{31}$PNMR (DMSO-$d_6$)

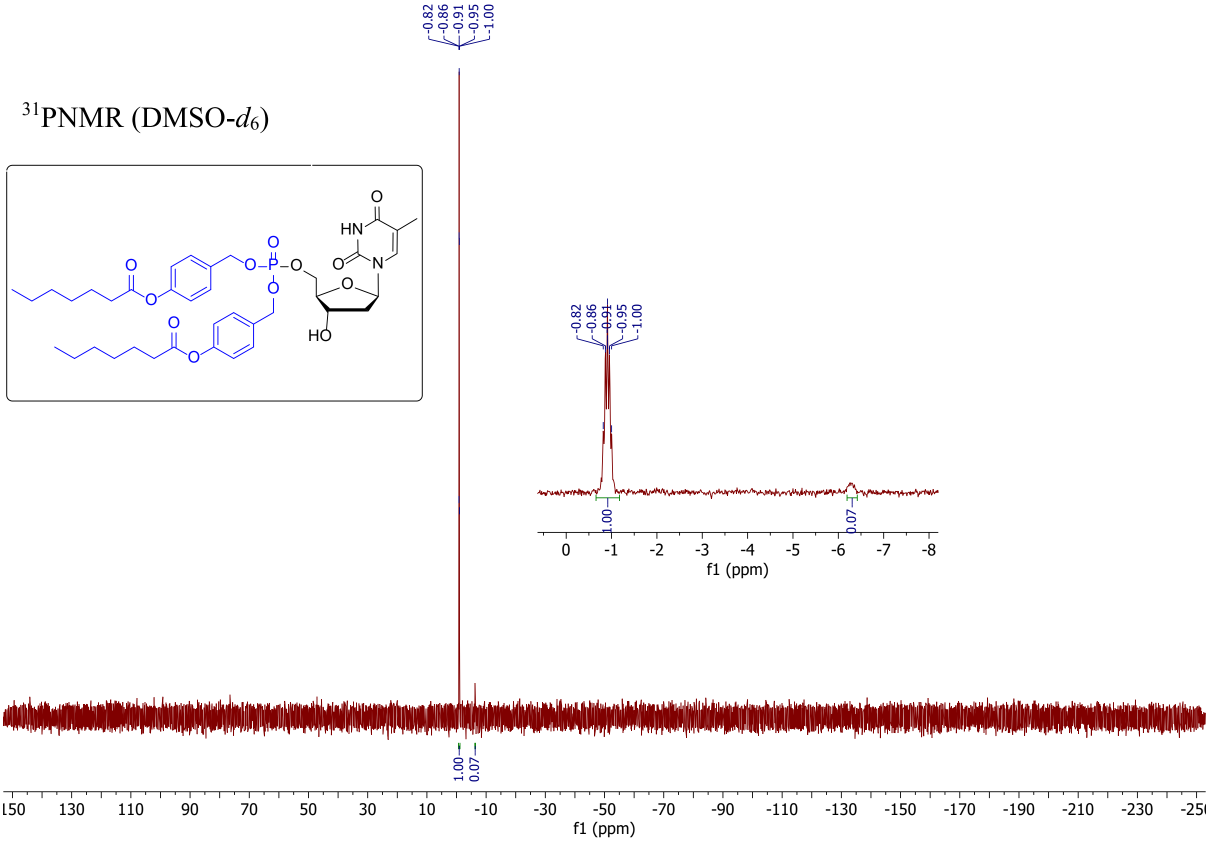

$^{31}$P{H}NMR (DMSO-$d_6$)

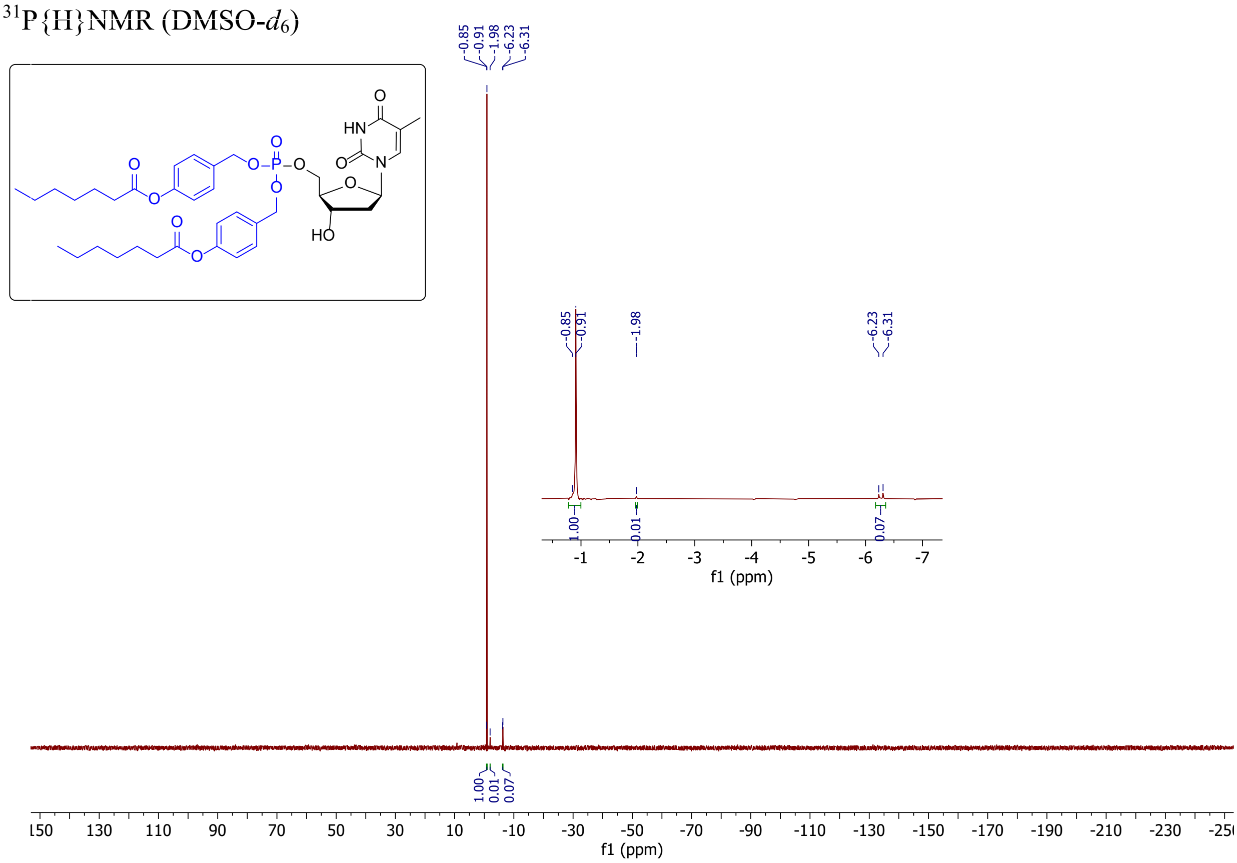

#### Supplementary references

Hilko, D. H., Fisher, G. M., Addison, R. S., Andrews, K. T., & Poulsen, S. A. (2023). Thymidine Kinase-Independent Click Chemistry DNADetect Probes for DNA Proliferation Assessment in Malaria Parasites. *ACS Chem Biol*, *18*(12), 2535-2543. https://doi.org/10.1021/acschembio.3c00530
